# Novel mouse models for perianal fistulizing Crohn’s disease reveal therapeutic value of interferon-γ antagonists

**DOI:** 10.64898/2026.06.04.730162

**Authors:** Xin Yao, Kaiming Ma, David H. Ballard, Echo Zhu, Xiuli Liu, Lingjie Huang, Chuwen Tian, James D. Quirk, Heather S. Ruiz, Tingyi Tan, Matthew A. Ciorba, Gwendalyn J. Randolph, Parakkal Deepak, Siyan Cao

## Abstract

**Background and Aims:** Perianal fistulizing Crohn’s disease (PFCD) is a challenging complication with poorly understood pathogenesis and limited treatment options, largely due to the lack of clinically relevant animal models. Interferon-gamma (IFN-γ) signaling is hyperactivated in human PFCD. We aimed to establish mouse models recapitulating human PFCD and to evaluate IFN-γ as a new therapeutic target.

**Methods:** Perianal fistulas were established in three mouse models with concurrent Crohn’s disease-like intestinal inflammation: wild-type (WT) mice with 2,4,6-trinitrobenzenesulfonic acid (TNBS)-induced proctocolitis, *Il10^−/−^*mice, and *TNF^Δ69AU/+^* mice. A modified MAGNIFI-CD index was developed for longitudinal fistula assessment in mice. Transcriptomic analysis and flow cytometry were conducted on mouse fistula tissue. Re-analysis of single-cell and spatial transcriptomics of human PFCD tissues was performed. Therapeutic benefits of anti-TNF-α, upadacitinib, and IFN-γ pathway antagonists were evaluated in the PFCD models.

**Results:** All three PFCD models sustained chronic perianal fistula tracts for at least 5 weeks after wire removal. All three models closely recapitulate the pathological and molecular features of PFCD in patients, as confirmed by clinical examination, MRI, histopathology, immunostaining, flow cytometry, and transcriptomics. IFN-γ signaling emerged as a central and conserved pathway across all three mouse models and human PFCD. Targeting of the IFN-γ pathway promptly improved fistula healing with mitigation of IFN-γ signaling, inflammation, and epithelial-to-mesenchymal transition (EMT). Moreover, combining IFN-γ and TNF-α blockade demonstrated augmented therapeutic efficacy compared to anti-TNF-α monotherapy.

**Conclusions:** These PFCD mouse models and imaging tools provide first reliable and clinically relevant platforms for mechanistic studies and therapeutic evaluation. IFN-γ signaling represents a potential therapeutic target warranting clinical investigation.

## Introduction

Perianal fistulizing Crohn’s disease (PFCD) affects up to 30% of CD patients^1^, and is associated with significant morbidity, impaired quality of life, therapeutic resistance, and escalating costs of healthcare. Despite the advent of anti-TNF-α biologics, only approximately 36% of patients achieve long-term complete response (defined as the absence of draining fistulas)^2^, and in severe refractory disease, 31%-49% ultimately require surgery^1^. The pathophysiology driving fistula formation and chronicity remain incompletely understood^3^, representing a critical barrier to the development of more effective therapies.

Accumulating evidence suggests that PFCD is not simply a consequence of local tissue injury, but rather a multifactorial process driven by persistent inflammation, epithelial barrier dysfunction, aberrant epithelial plasticity, and dysregulated immune-stromal interactions. Current models propose that defects in the intestinal epithelial barrier permit infiltration of pathogen-associated molecular patterns (PAMPs) into the intestinal mucosa, triggering chronic inflammatory responses and sustained cytokine production. Persistent inflammatory cytokine signaling, particularly involving TNF-α, TGF-β, IL-13, and related pathways, is believed to sustain EMT activation and promote chronic nonhealing fistula tracts^3, 4^. Our recent single-cell and spatial transcriptomic analyses further revealed hyperactivated IFN-γ signaling in both the fistula tract in PFCD patients compared to idiopathic fistula, together with increased IFN-γ signaling in rectocolonic and ileal mucosa of CD patients with perianal fistulas compared to those without fistulizing disease. Interestingly, ligand-receptor interaction analysis also showed upregulation of MMP9-CD44 and VIM-CD44 interactions^5^, suggesting a potential link between IFN-γ signaling and EMT, which has been identified as a key driver of fistula tract formation and persistence^4, 6, 7^.

Mechanistic studies and drug discovery for PFCD have been markedly hampered by the lack of reliable and clinically relevant preclinical models^8–12^. Mouse models offer critical advantages over larger animals, including low cost, ease of use, and amenability to genetic manipulation, yet no reliable mouse model has been available^3, 13^. Moreover, no standardized imaging-based outcome measure exists for longitudinal fistula monitoring in preclinical animals, further constraining therapeutic studies^9, 10^.

Here, we established three models of PFCD that closely resemble human PFCD based on clinical examination, MRI, histopathology, cellular and molecular analyses, and therapeutic response. We further demonstrated that heightened IFN-γ signaling is a conserved molecular feature across our mouse models and human PFCD. Inhibition of IFN-γ signaling promoted fistula healing, and combined IFN-γ and TNF-α blockade showed superior efficacy compared to anti-TNF monotherapy. Together, these findings provide mechanistic insight into PFCD pathogenesis, establish clinically relevant mouse models for mechanistic studies and drug discovery, and support dual targeting of IFN-γ and TNF-α as a potential therapeutic strategy for refractory PFCD.

## Methods

### Establishing PFCD models in mice

Three mouse models of CD were used to create PFCD models in this study: wild-type (WT) mice with 2,4,6-trinitrobenzenesulfonic acid (TNBS)-induced proctocolitis^14^, *Il10^−/−^*mice^15^ and *TNF^Δ69AU/+^* mice with continuous, low-level overproduction of TNF-α^16^. A standardized mechanical procedure was established to induce perianal fistula in the three models as well as for non-IBD fistula in WT mice. Mice were anesthetized with 2.5% isoflurane and placed in the supine position. A 16-gauge guide needle was used to create a tract from the rectal lumen to the perianal skin at the 10 o’clock position, with the external opening located approximately 0.5 cm from the anus. Then, a bundle of five precut, autoclaved stainless-steel wires (1 mm in diameter) was introduced through the needle lumen. After needle withdrawal, the wire bundle was left in place and shaped into a ring to keep the fistula tract open. For the WT-TNBS model, TNBS (Sigma, Catalog #P2297) was administered intrarectally on day 5 after fistula creation and continued twice weekly throughout the entire experimental period, including post-wire removal phases. TNBS stock solution at 5% (w/v) in H_2_O was diluted 1:1 with 100% ethanol to generate 2.5% (w/v) TNBS in 50% ethanol immediately before use. A volume of 100 µL solution was administrated intrarectally to a depth of approximately 1 cm from the anus in each mouse. For non-IBD fistula, WT mice did not receive TNBS after fistula creation. All mice were monitored for body weight change and clinical signs including diarrhea, hematochezia, and reduced activity during the experiment.

All mice used for experiments were on a C57BL/6 background and 8-12 weeks of age at the time of experiments. No sex-based differences were observed; therefore, both sexes were included in the study. Mice were housed in regular filter-top cages with free access to sterile water and food in a pathogen-free barrier facility fully staffed and equipped by the Washington University Division of Comparative Medicine, with daily monitoring by highly trained staff. Environmental conditions were reproducibly regulated for temperature, humidity, lighting, caging, bedding, and water.

## Results

### Novel PFCD mouse models maintain chronic inflamed perianal fistula tract

To establish reproducible and clinically relevant mouse models of PFCD, a standardized mechanical procedure was used to induce perianal fistulas in three CD models: WT-TNBS, *Il10^−/−^*, and *TNF^Δ69AU/+^*mice (Supplemental Figure 1*A*). The perianal wire was kept in place for 35 days to allow fistula tract maturation, confirmed by physical examination and probe exploration of fistula tract patency (Figure 1*A*). After wire removal, fistulas were evaluated by weekly MRI for at least five weeks. A subset of mice underwent longitudinal MRI monitoring every other week until week 9. MRI is an essential tool for diagnosis and longitudinal monitoring of CD perianal fistulas in clinical practice. To further enable standardized and quantitative longitudinal monitoring, we adapted the MAGNIFI-CD scoring system for human PFCD^17^ to mouse settings, integrating MRI-based assessment of fistula tract activity, perifistular tissue changes, and rectal wall inflammation to better mimic the clinical situation (Supplemental Table 1).

**Figure 1.**
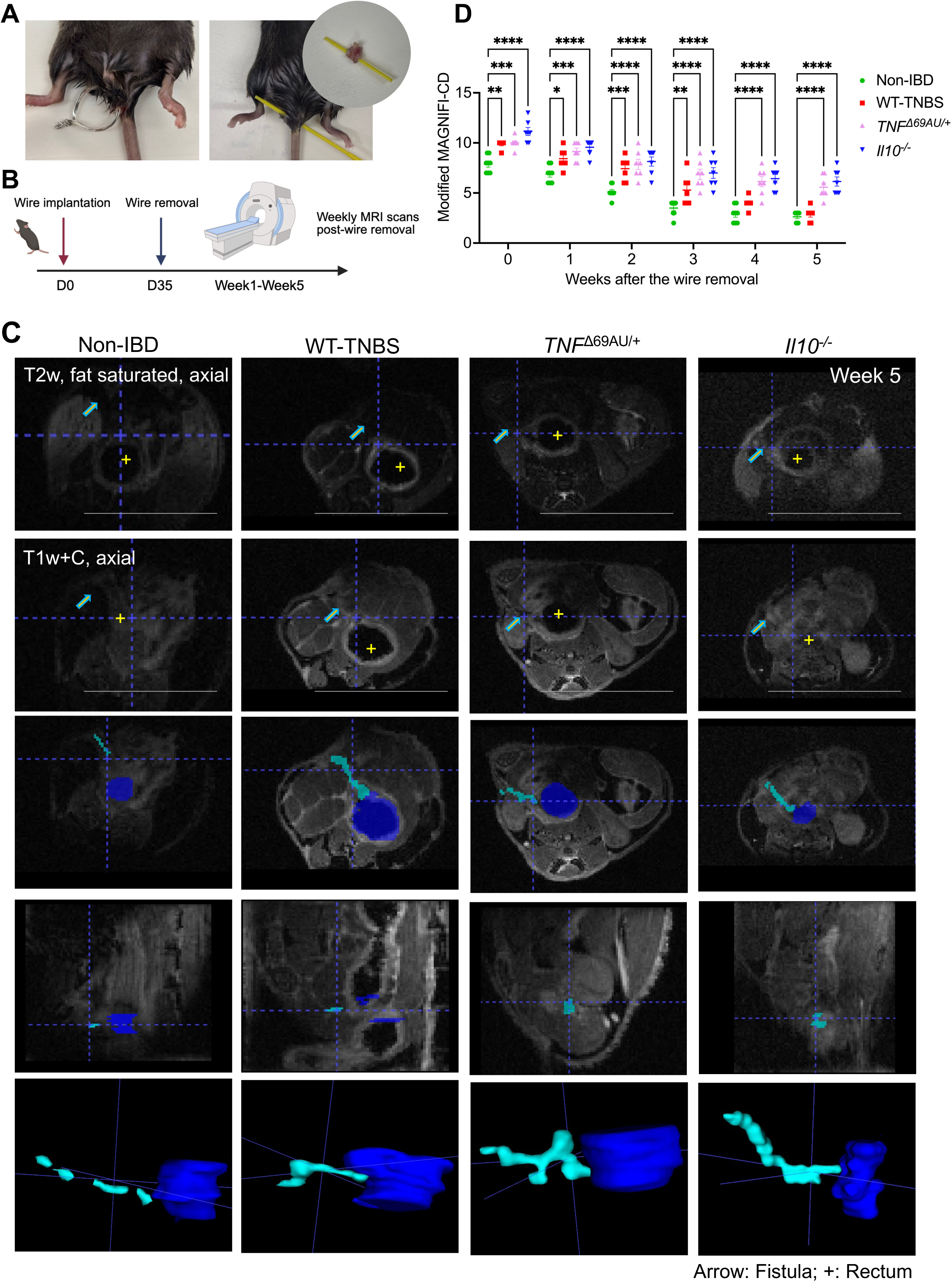
Perianal fistula induction in mouse models and longitudinal MRI assessment. (*A*) Macroscopic features of perianal fistula in the models: perianal wire implantation with the wire in situ (left), exploration of the fistula with a probe demonstrating a fistulous tract from the anus to the perianal skin after fistula maturation on Day 35 (middle), and the fistula tract after dissection (right). (*B*) Schematic overview of the experimental design: perianal wire was implanted on Day 0 and removed on Day 35, followed by weekly MRI scans from Week 0 to Week 5 post-wire removal (the MRI was continued until Week 9 in a subset of mice). (*C*) Representative multiparametric MRI images at Week 5 in PFCD models (WT-TNBS, *Il10^−/−^* and *TNF^Δ69AU/+^*) and non-IBD fistula controls. T2-weighted fat-saturated and T1-weighted with contrast (T1w+C) axial images with corresponding segmented views and 3D reconstructions of fistula tracts (cyan) and rectum (blue). Arrow: fistula tract; +: rectum. Scale bars, 1cm. (*D*) Longitudinal modified MAGNIFI-CD scores from Week 0 to Week 5 post-wire removal in PFCD models and non-IBD controls (n=6-9). Data are presented as mean ± SEM. Statistical comparisons were performed using two-way ANOVA. Experiments were repeated at least 3 times. \**p*<0.05, \*\**p*<0.01, \*\*\**p*<0.001, \*\*\*\**p*<0.0001.

All three PFCD models sustained patent, chronic fistula tracts connecting the rectum and perianal area for at least five weeks, as confirmed by both physical examination and multiparametric MRI (Figure 1*B*-*C*). T2-weighted fat-saturated and T1-weighted with contrast axial images demonstrated persistent fistula tracts with surrounding inflammation, while 3D reconstructions revealed complex branching morphology in all three PFCD models, resembling findings in PFCD patients^4^ (Figure 1*C*). At week 9, perianal fistula tracts remained visible on MRI in *Il10^−/−^*and *TNF^Δ69AU/+^* mice, with sustained perifistular and rectum inflammation (Supplemental Figure 1*B*). In contrast to PFCD models, non-IBD fistula controls (WT mice without TNBS-induced inflammation) showed progressive fistula healing over the follow-up period, with >50% tract hypointensity on T2 imaging consistent with scarring by Week 5. Significantly elevated MAGNIFI-CD scores were observed in all three PFCD models compared to non-IBD controls throughout the entire follow-up period (Figure 1*D*), demonstrating that our models recapitulate the chronic, active fistula disease course in PFCD patients.

### PFCD mouse models recapitulate histological and molecular features of human PFCD

In addition to examination and MRI, histopathology of perianal specimens confirmed inflammatory fistula tracts connecting the rectum and perianal skin in all three PFCD models, with elevated acute and chronic inflammatory infiltration and granulation tissue formation on H&E staining (Figure 2*A*). Histological scores for both fistula tract inflammation and proctitis were significantly higher in all three PFCD models compared to non-IBD controls (Figure 2*B*, Supplemental Figure 2).

**Figure 2.**
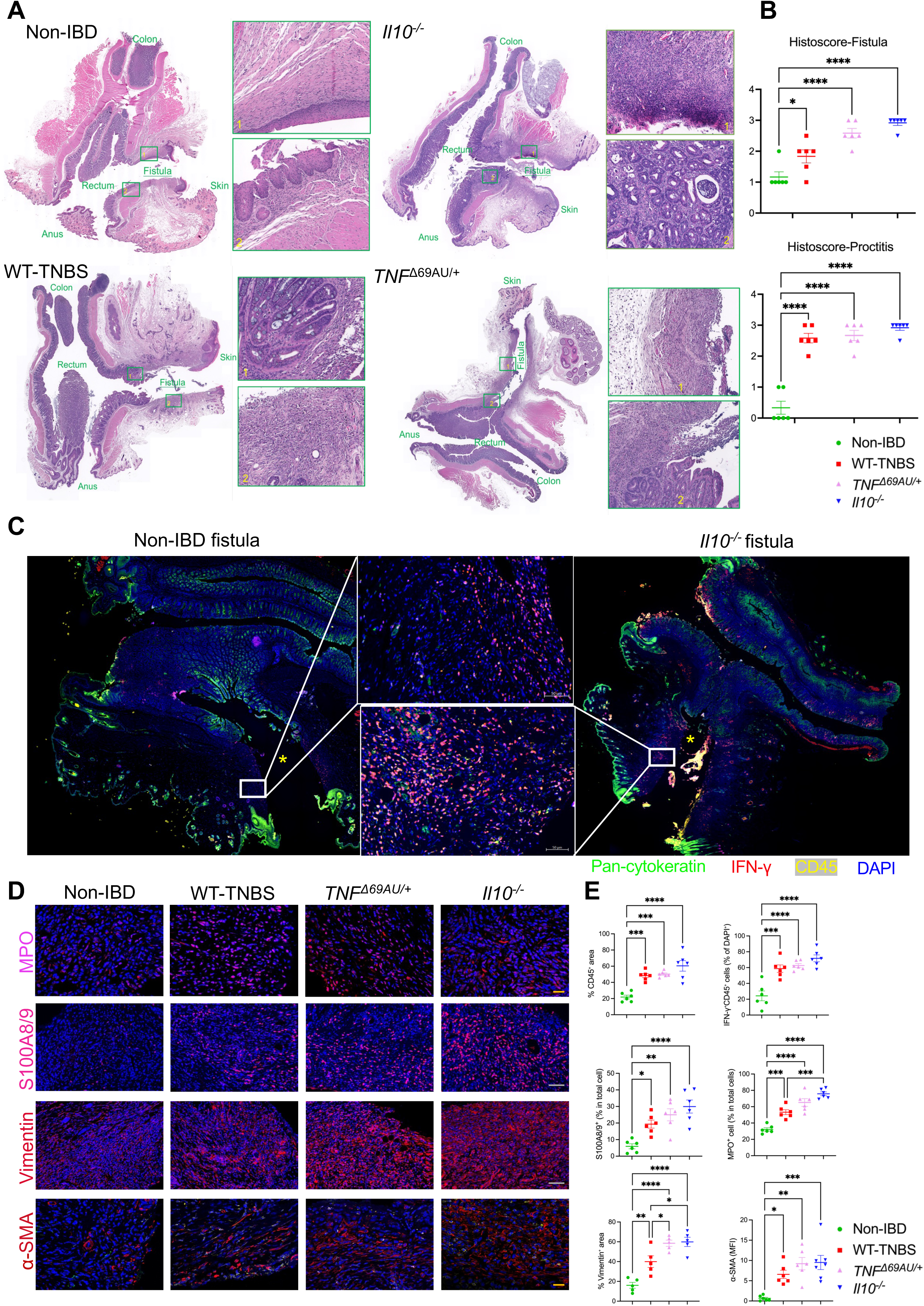
Histological and immunofluorescence characterization of perianal fistula tracts in PFCD mouse models. (*A*) Representative H&E staining showing perianal fistula connecting rectum and perianal skin in non-IBD fistula controls (surgical perianal fistula in WT mice without TNBS treatment) and PFCD models (WT-TNBS, *Il10^−/−^*and *TNF^Δ69AU/+^*). Insets: higher magnification of the fistula tract and surrounding tissue. (*B*) Histological scores for fistula tract inflammation (Histoscore-Fistula) and proctitis (Histoscore-Proctitis) in PFCD models and non-IBD controls (n=5-7). (*C*) Representative immunofluorescence images of fistula tracts from non-IBD controls and *Il10^−/−^*model stained for pan-cytokeratin (green), IFN-γ (red), CD45 (yellow), and DAPI (blue). Inset: higher magnification of the boxed region. Yellow asterisk indicates the fistula tract. (*D*) Representative immunofluorescence images of fistula tracts from PFCD models and non-IBD controls stained for MPO, S100A8/9, Vimentin, and α-SMA, with DAPI counterstain (blue). Scale bars, 20 µm (MPO, α-SMA) and 50 µm (S100A8/9, Vimentin). (*E*) Quantification of immunofluorescence signals in (*C*) and (*D*) across PFCD models and non-IBD controls (n=6). MFI, mean fluorescence intensity. Data are presented as mean ± SEM. Statistical comparisons were performed using one-way ANOVA with Tukey’s post-hoc test. Experiments were repeated at least 3 times. \**p*<0.05, \*\**p*<0.01, \*\*\**p*<0.001, \*\*\*\**p*<0.0001.

As our recent single-cell and spatial transcriptomic analyses of human PFCD tissue revealed hyperactivated IFN-γ signaling in fistula tracts^5^, we next examined whether this pathway is similarly activated in PFCD mouse models. IFN-γ^+^ cells were markedly enriched in the fistula tissue of PFCD mice compared to non-IBD controls, with a notable increase in CD45^+^IFN-γ^+^ immune cells by immunofluorescence (Figure 2*C-E*). Inflammatory markers MPO and S100A8/9 and tissue remodeling markers vimentin and α-SMA were similarly increased across all PFCD mice (Figure 2*D-E*), collectively demonstrating that the PFCD mouse models exhibit the key histological and molecular features of human PFCD. To further characterize the PFCD models, we performed bulk RNA sequencing of perianal fistula tissues after wire removal. Principal component analysis (PCA) demonstrated clear transcriptomic separation between PFCD models and non-IBD controls, indicating distinct transcriptomic profiles in PFCD fistula tissues (Figure 3*A*). Venn diagram analysis of differentially expressed genes identified 1,283 commonly dysregulated genes shared across all three PFCD models versus non-IBD controls (Figure 3*A*), suggesting shared disease mechanisms across different models. GSEA revealed consistent upregulation of IFN-γ response, inflammation, EMT, fibrosis, IL-17 signaling, TL1A-DR3 pathway, and TNF superfamily pathways, as well as downregulation of adaptive cellular response including chaperones, autophagy, epithelial integrity programs including tight junction, and keratinization across the PFCD models (Figure 3*B*, *C*). Specifically, all PFCD fistulas showed elevations in EMT genes including *Vim*, *Snai1*, *Snai2*, *Twist1*, *Zeb2*, and *Mmp9*, fibrosis-associated genes including *Postn*, *Pdgfra*, *Lox* (Supplemental Figure 3*A*), as well as TNF and TNF receptor superfamily members (Supplemental Figure 3*B-C*). Given that TL1A (encoded by *Tnfsf15*) has emerged as a promising therapeutic target in IBD and reported to be elevated in rectum tissue from PFCD patients^5^, we next examined the expression of TL1A and its receptor DR3 (encoded by *Tnfrsf25*) in the models. Normalized expression of DR3 (*Tnfrsf25*) was significantly elevated in *TNF^Δ69AU/+^*(*p* = 0.00629) and *Il10^−/−^* (*p* = 0.002) fistulas, with a trend also observed in WT-TNBS fistulas (*p* = 0.0665). Similarly, TL1A (*Tnfsf15*) expression was significantly increased in *Il10^−/−^*(*p* = 0.00719) fistulas, with a trend in WT-TNBS (*p* = 0.0831) and *TNF^Δ69AU/+^* (*p* = 0.0587) mice (Supplemental Figure 3*D*). Examination of downstream TL1A-DR3 pathway components revealed that *Mapk8*, encoding JNK1, was notably and consistently upregulated across all PFCD models (Supplemental Figure 3*E*).

**Figure 3.**
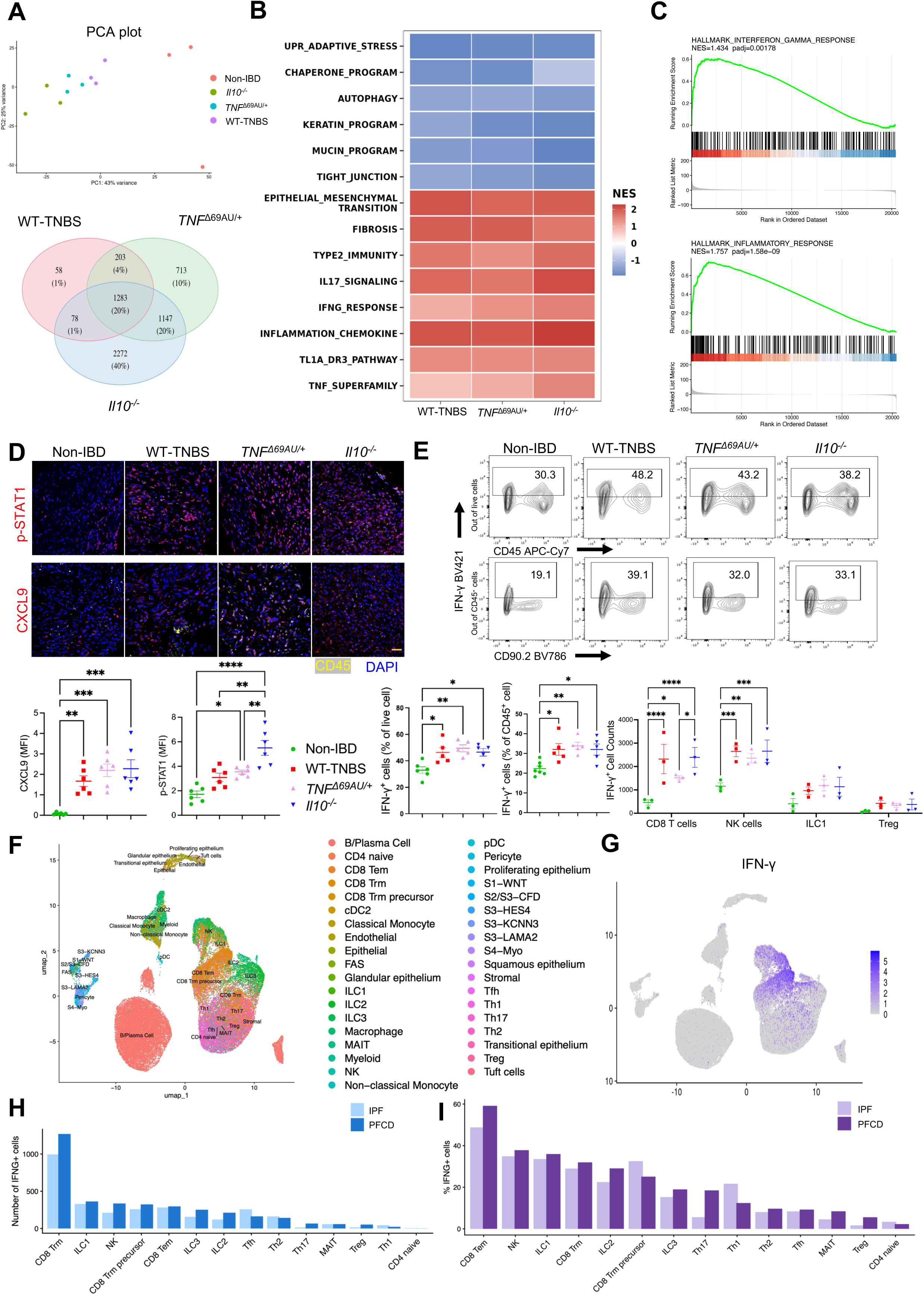
Transcriptomics of perianal fistula tissues in PFCD mouse models and human PFCD reveals shared pathways. (*A*) PCA plot of RNA-seq data from fistula tissues of PFCD models and non-IBD controls (upper). Venn diagram of differentially expressed genes (DEGs) across the three PFCD models and non-IBD fistula controls (lower). (*B*) Heatmap of normalized enrichment scores (NES) from GSEA across the three PFCD models. (*C*) GSEA plots of IFN-γ response and inflammatory response gene sets in PFCD fistula tissue versus non-IBD controls. (*D*) Representative immunofluorescence images and quantification of IFN-γ downstream targets p-STAT1 and CXCL9 in fistula tracts across PFCD models and non-IBD controls (n=6). MFI, mean fluorescence intensity. Scale bars, 20 µm. (*E*) Representative flow cytometry plots and quantification of IFN-γ+ cells across immune cell subsets in fistula tissues of PFCD models and non-IBD controls (n=3-6). Experiments were repeated at least 3 times. (*F*) UMAP of scRNA-seq data from human IPF and PFCD tissues with cell cluster annotation. (*G*) UMAP projection showing the distribution of IFN-γ expression across distinct cell clusters. Color intensity from light gray to deep blue indicates increasing expression levels. Number (*H*) and percentage (*I*) of IFN-γ^+^ cells across immune subpopulations in IPF and PFCD tissues. Data are presented as mean ± SEM. Statistical comparisons were performed using one-way ANOVA with Tukey’s post-hoc test. \**p*<0.05, \*\**p*<0.01, \*\*\**p*<0.001, \*\*\*\**p*<0.0001.

Given the emerging evidence implicating gut microbial dysbiosis in the pathogenesis of PFCD^3^, we performed 16S rRNA gene sequencing on fecal samples from WT mice with non-IBD fistula, WT-TNBS and *Il10^−/−^* models. Both PFCD models exhibited reduced alpha diversity and distinct shifts in overall microbial community composition relative to controls in PCoA analysis (Supplemental Figure 4*A, B*). At the genus level, a consistent pattern of dysbiosis emerged across both models: depletion of potentially protective taxa including *Alistipes* and *Paramuribaculum*, alongside marked enrichment of *Muribaculum*^18–20^ (Supplemental Figure 4*C, D*).

### IFN-γ pathway is activated in PFCD fistula tissues across mouse models and patients

Activation of IFN-γ downstream signaling was further confirmed at the protein level by immunofluorescence, demonstrating significantly elevated p-STAT1 and CXCL9 expression in fistula tracts of all three PFCD models versus non-IBD controls (Figure 3*D*). Flow cytometry further detected expanded IFN-γ^+^ immune cells in PFCD fistula tissues, with CD8^+^ T cells and NK cells showing the most prominent increases among CD45^+^ immune cells (Figure 3*E*).

To validate the clinical relevance of these findings in mice, we re-analyzed scRNA-seq data from human PFCD and idiopathic fistula (IPF) tissues^5^. UMAP visualization demonstrated that IFNG expression was predominantly detected in T cells and innate lymphoid cells (ILCs) (Figure 3*F-G*). Subpopulation analysis revealed that IFN-γ^+^ cells were elevated across multiple immune subsets in PFCD versus IPF specimens, with the most prominent increases observed in CD8^+^ T cell subsets, NK cells, and ILC1s (Figure 3*H-I*). To delineate the cellular targets of IFN-γ signaling within the fistula microenvironment, we performed cell-cell communication analysis using human PFCD scRNA-seq data^5^. IFN-γ-producing immune cells are predominantly CD8^+^ T cells, ILC1s, and NK cells. They similarly exhibited markedly increased communication to myeloid, stromal, and epithelial cells in PFCD compared to IPF (Figure 4*A*), revealing broadly activated IFN-γ signaling across multiple tissue compartments within the fistula niche. Among the strongest recipient cell populations, macrophages emerged as a key downstream target. Both M1 pro-inflammatory and M2 tissue-remodeling scores were significantly upregulated in PFCD macrophages compared to their IPF counterparts (Supplemental Figure 5*A*), indicating that macrophages within the fistula niche adopt a dual-activated state that may simultaneously drive inflammation and aberrant tissue remodeling.

**Figure 4.**
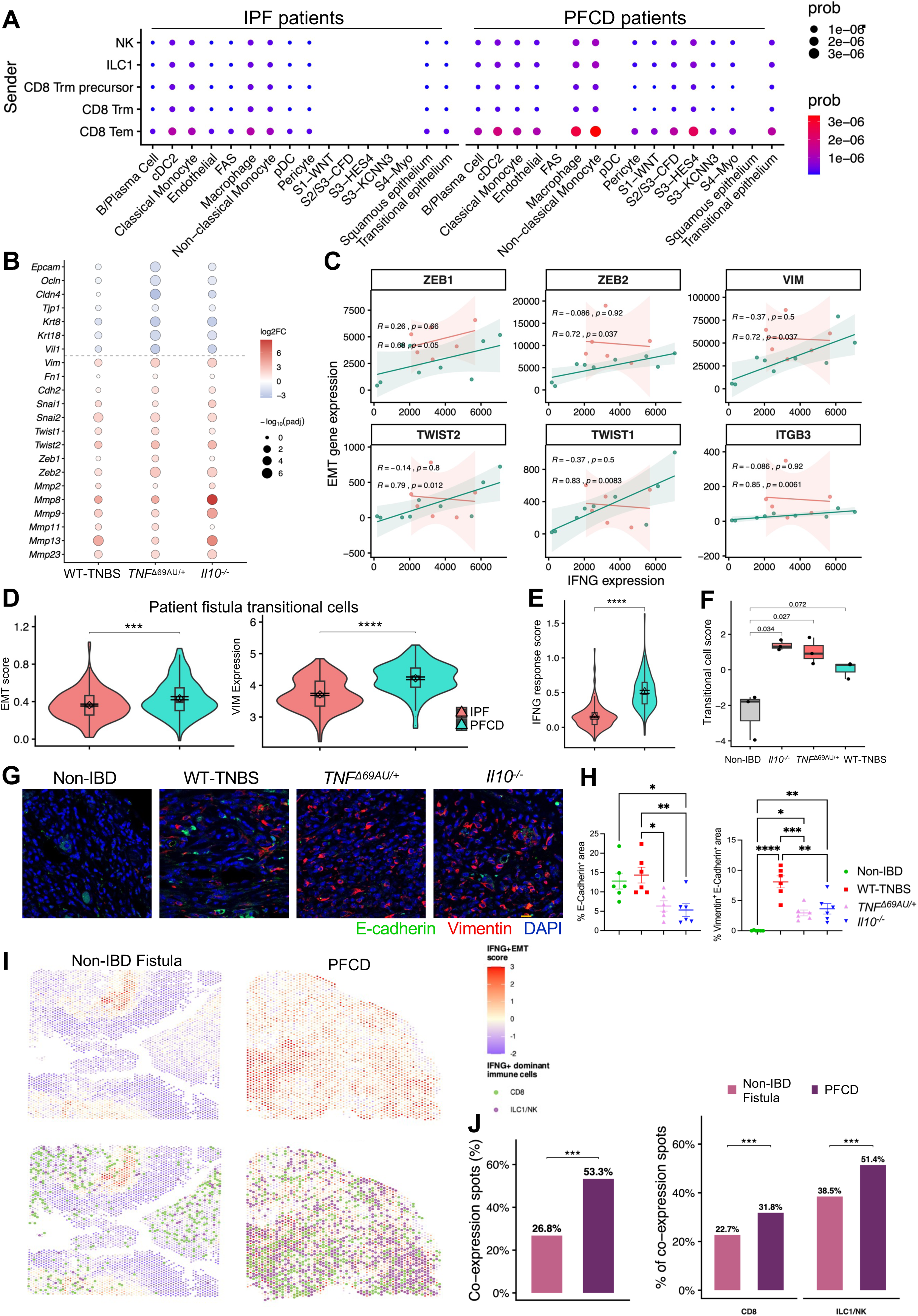
Shared IFN-γ and EMT pathways in the fistula tissues of PFCD mouse models and patients. (*A*) Bubble plot indicating cell-cell communication probabilities between immune cells (Senders) and stromal/epithelial compartments (Receivers) in human IPF and PFCD fistula tissues. (*B*) Bubble plot of log2 fold change and statistical significance of EMT and tissue remodeling-related genes across PFCD mouse models compared to non-IBD controls. (*C*) Scatter plots depicting Spearman correlations between IFNG expression and the expression of EMT marker genes (*ZEB1*, *ZEB2*, *VIM*, *TWIST2*, *TWIST1*, and *ITGB3*) in the fistulas of IPF (pink) and PFCD (green) patient cohorts. Spearman correlation coefficients (R) and corresponding p-values are shown for each group. Shaded areas represent 95% confidence intervals. (*D*) Violin plots comparing EMT scores (left) and VIM expression (right) in fistula transitional cells from IPF and PFCD patients. (*E*) Violin plots depicting IFNG pathway response score in fistula transitional cells from IPF and PFCD patients. (*F*) Box plots comparing fistula transitional cell score in PFCD mouse models and non-IBD controls. (*G*) Representative immunofluorescence images of fistula tract from PFCD mouse models and non-IBD controls, stained with DAPI (blue), E-cadherin (green), and Vimentin (red). Scale bars, 20 µm. (*H*) Quantification of *G* for E-cadherin⁺ area and Vimentin⁺ E-cadherin⁺ double-positive area. Data are presented as mean ± SEM. Statistical significance was determined by one-way ANOVA with multiple comparisons. (*I*) Spatial transcriptomic analysis of fistula tissue sections from PFCD and non-IBD control (diverticular fistula) patients. Upper: combined IFN-γ and EMT scores, defined as the sum of z-scaled IFN-γ and EMT module scores, visualized across all tissue spots. Lower: spatial distribution of major IFN-γ⁺ immune cells (CD8⁺ T cells and ILC1/NK cells), defined as spots with module score > 0.1 for the respective cell type signature, overlaid on the combined IFN-γ/EMT score. (*J*) Bar plot showing the percentage of IFN-γ/EMT co-expressing spots (defined as spots with both IFN-γ and EMT module scores > 0) and the percentage of IFN-γ/EMT co-expression spots attributed by CD8^+^ T cells and ILC1/NK cells in PFCD versus non-IBD controls (diverticular fistula) patients (From the published spatial transcriptomic dataset; McGregor, 2026.) \**p*<0.05, \*\**p*<0.01, \*\*\**p*<0.001, \*\*\*\**p*<0.0001.

### IFN-γ signaling correlates with EMT in PFCD fistula tissues

EMT and tissue remodeling are recognized as hallmarks in human PFCD^3, 5, 21^. To assess whether these processes are consistently activated across experimental models, we examined the expression of canonical EMT-associated genes in all three PFCD mouse models. Key mesenchymal markers were consistently and significantly upregulated across all three PFCD models compared to non-IBD controls. Conversely, genes regulating epithelial integrity and keratinization, including *Cdh1*, *Epcam*, *Ocln*, *Cldn4*, *Tjp1*, *Krt8*, and *Krt18*, were markedly downregulated, indicative of epithelial barrier disruption (Figure 4*B*). Together, these findings demonstrate that EMT and tissue remodeling are robustly engaged across PFCD mouse models. Similarly, re-analysis of our patient single cell datasets^5^ uncovered that epithelial cells from PFCD fistulas exhibited augmented expression of mesenchymal markers *ZEB1* and *VIM*, indicating an ongoing EMT process (Supplemental Figure 5*B*).

The simultaneous upregulation of both IFN-γ response and EMT pathways across PFCD models prompted us to investigate whether these two programs are functionally linked. In human PFCD fistulas, IFNG expression showed strong and significant positive Spearman correlations with a broad set of EMT-related genes (Figure 4*C*, Supplemental Figure 5*C*). Notably, these correlations were largely absent or substantially weaker in IPF tissues, suggesting that the relationship between IFN-γ signaling and EMT gene programs are preferentially enriched in the PFCD fistula microenvironment.

To determine which specific epithelial cell subsets are undergoing EMT-like reprogramming in human PFCD fistulas, we examined the expression of key EMT markers VIM and S100A4, as well as the overall EMT score, across different epithelial subsets in our scRNA-seq dataset^5^. Transitional cells, which have been reported to occupy an intermediate state between epithelial and stromal cells, expressing both epithelial markers (KRT8 and KRT18) and stromal cell markers, were identified in human PFCD fistulas (Figure 4*D*, Supplemental Figure 5*D*). Specifically, transitional epithelial cells in PFCD exhibited significantly upregulated IFN-γ response scores compared to IPF (Figure 4*E*). Given that TNF and TGF-β are also recognized as potential inducers of EMT in PFCD, we further investigated whether the EMT program observed here was specific to IFN-γ signaling. Neither the TNF nor the TGF-β pathway showed significant upregulation in PFCD versus IPF (Supplemental Figure 5*E*). These findings highlight the vital and disease-specific role of IFN-γ in driving EMT in human PFCD. To validate the above findings in the PFCD models, we assessed transitional cell scores in mouse fistula tissues. We combined the EMT score and epithelial score to evaluate the transitional state. Consistently, transitional cell scores were augmented in PFCD models versus non-IBD controls (Figure 4*F*). To corroborate the EMT process in PFCD models, we performed immunofluorescence staining for E-cadherin and Vimentin of perianal tissues (Figure 4*G*). E-cadherin⁺ area, reflecting epithelial content, was higher in non-IBD and WT-TNBS groups compared to *Il10^−/−^* and *TNF*^Δ69AU/+^ models; while the proportion of Vimentin⁺E-cadherin⁺ double-positive cells, representing cells in an intermediate EMT state, was significantly increased across all three PFCD models (Figure 4*H*).

Fistula-associated stromal (FAS) fibroblasts have been identified as the predominant and functionally distinct stromal cell population within CD fistula tracts^22^. To examine whether IFN-γ signaling is specifically enriched in this stromal cell subset, we analyzed IFN-γ, TNF, and TGF-β response scores in FAS fibroblasts from PFCD and IPF^5, 22^. IFNG response scores were significantly increased in FAS fibroblasts from PFCD compared to IPF, while TNF and TGF-β response scores showed no significant difference between groups (Supplemental Figure 6*A*). Furthermore, FAS fibroblasts exhibited significantly higher IFN-γ response scores compared to non-FAS stromal cells (Supplemental Figure 6*B*). The data suggest that IFN-γ acts on stromal cell compartments in shaping PFCD microenvironment.

### Spatial transcriptomic analysis reveals co-localization of IFN-γ signaling and EMT within the PFCD fistula tissue microenvironment

To spatially resolve the relationship between IFN-γ signaling and EMT within the intact fistula tissue architecture, we reanalyzed spatial transcriptomic data using Visium datasets from PFCD patients^22^. Combined IFN-γ and EMT scoring uncovered markedly elevated co-localization of these two signals in representative PFCD versus non-IBD control (diverticular fistula) specimens (Figure 4*I*). Further integration of immune cell spatial signatures revealed that IFN-γ-producing immune cells, including CD8⁺ T cells and ILC1/NK cells, were preferentially enriched within regions of high IFN-γ and EMT co-expression in PFCD (Figure 4*I*). In contrast, this coordinated spatial relationship was largely absent in non-IBD control specimens, suggesting that the IFN-γ to EMT spatial axis is a disease-specific feature of PFCD. Consistent with these findings, analysis of an independent spatial transcriptomic dataset from Cao et al.^5^ confirmed similar spatial co-enrichment of IFN-γ signaling, EMT score, and IFN-γ⁺ immune cells in PFCD fistula tissues (Supplemental Figure 7*A-B*). Furthermore, we quantitatively validated these spatial findings using a generalized linear mixed model. The proportion of IFN-γ/EMT co-expressing spots (defined as spots with both IFN-γ and EMT scores > 0) was significantly higher in PFCD fistulas compared to non-IBD fistulas (53.3% vs. 26.8%, *p*<0.001). Consistently, the co-localization of IFN-γ-producing immune cells with IFN-γ/EMT co-expressing spots was also significantly enriched in PFCD versus non-IBD fistulas, with co-expression spot proportions of 31.8% vs. 22.7% (*p*<0.001) for CD8⁺ T cells and 51.4% vs. 38.5% (*p*<0.001) for ILC1/NK cells (Figure 4*J*).

### The PFCD models are therapeutically responsive to anti-TNF and benefit from IFN-γ pathway antagonists

To determine the therapeutic responsiveness of our PFCD models, we first tested anti-TNF antibody treatment, a first-line biologic therapy for PFCD patients. As *TNF^Δ69AU/+^* and *Il10^−/−^*mice share comparable imaging, histopathological, molecular, and transcriptomic features, and given the broader applicability of *Il10^−/−^* mice as an established CD model, we selected WT-TNBS and *Il10^−/−^* models for therapeutic validation. Anti-TNF treatment (0.5mg/mouse, twice weekly) significantly reduced modified MAGNIFI-CD scores compared to PBS controls (Figure 5*A, B*), confirming that our models are therapeutically responsive.

**Figure 5.**
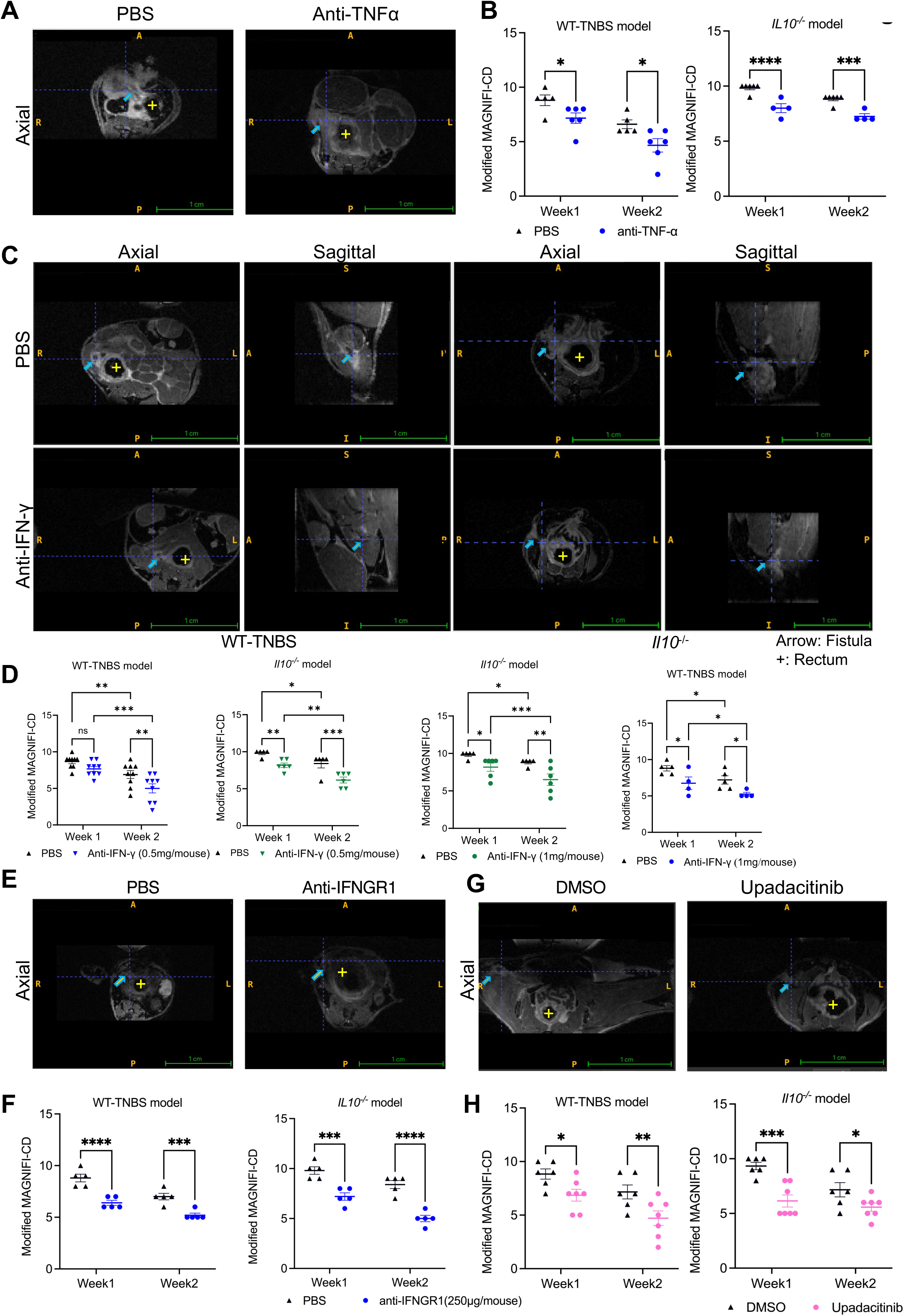
PFCD mouse models benefit from anti-TNF and IFN-γ pathway antagonists. (*A*) Representative MRI images at Week 1 and 2 following anti-TNF antibody or PBS treatment in WT-TNBS and *Il10^−/−^* models (n=6). (*B*) Modified MAGNIFI-CD scores at Week 1 and 2 following anti-TNF antibody or PBS treatment in WT-TNBS and *Il10^−/−^* models (n=6). (*C*) Representative axial and sagittal MRI images of PBS and anti-IFN-γ treated mice in WT-TNBS and *Il10^−/−^* models. Arrow: fistula tract; +: rectum. (*D*) Modified MAGNIFI-CD scores at Week 1 and 2 following anti-IFN-γ antibody (0.5mg/mouse or 1mg/mouse) or PBS treatment in WT-TNBS and *Il10^−/−^*models (n=4-9). (*E*) Representative axial MRI images of anti-IFNGR1 or PBS treated mice. (*F*) Modified MAGNIFI-CD scores at Week 1 and 2 following anti-IFNGR1 antibody (250μg/mouse) or PBS treatment in WT-TNBS and *Il10^−/−^*models (n=5). (*G*) Representative axial MRI images of upadacitinib or DMSO treated mice. (*H*) Modified MAGNIFI-CD scores at Week 1 and 2 following upadacitinib or DMSO treatment in WT-TNBS and *Il10^−/−^* models (n=6-7). Data are presented as mean ± SEM. Statistical comparisons were performed using unpaired t-test. Experiments were repeated at least 3 times. \**p*<0.05, \*\**p*<0.01, \*\*\**p*<0.001, \*\*\*\**p*<0.0001.

Building on the consistent activation of IFN-γ signaling across PFCD models and patients, we next directly inhibited this pathway using anti-IFN-γ antibody at two doses (0.5mg or 1mg/mouse, twice weekly). A reduction in MAGNIFI-CD scores was observed in both WT-TNBS and *Il10^−/−^*models at both doses, with MRI demonstrating reduced fistula tract inflammation and perianal tissue changes compared to untreated controls (Figure 5*C, D*). IFN-γ signals through IFNGR to activate JAK-STATs and drive interferon-stimulated gene (ISG) transcription^23^. Indeed, blockade of IFNGR1 (anti-IFNGR1, 0.25 mg/mouse, twice weekly) and JAK1 inhibition (upadacitinib, 20 mg/kg, every other day) both reduced MAGNIFI-CD scores in PFCD models (Figure 5*E–H*), collectively demonstrating that IFN-γ pathway activation plays an essential role in PFCD pathogenesis.

To characterize the molecular effects of IFN-γ pathway blockade, bulk RNA-seq of fistula tissues from anti-IFN-γ-treated and untreated WT-TNBS model revealed broad transcriptomic reversal of the key molecular features identified in PFCD fistula tissue (Figure 6*A*). Anti-IFN-γ treatment resulted in substantial downregulation of canonical IFN-γ signaling genes including *Stat1*, *Irf1*, and *Il1b*, as well as ISG genes *Ifit1/2/3*, confirming effective on-target suppression of IFN-γ pathways in the fistula tissues. Notably, immune suppression markers *Cd274* (encoding PD-L1) and *Ido1*, which are known downstream targets of IFN-γ and mediators of immune evasion, were also significantly reduced following treatment (Supplemental Figure 8*A*). In contrast, genes for epithelial integrity and repair including *Ocln*, *Muc1*, and *Muc13* were significantly induced in anti-IFN-γ-treated fistula tissues compared to untreated controls (Supplemental Figure 8*B*), suggesting that blockade of IFN-γ signaling is associated with restoration of mucosal barrier function. In addition, EMT-related genes including *Vim* and *S100a4*, as well as matrix-degrading enzyme genes *Mmp7* and *Mmp8*, were all suppressed in anti-IFN-γ-treated fistula tissues (Figure 6*B*), providing direct evidence that IFN-γ signaling promotes EMT and pathogenic tissue remodeling programs in PFCD. GSEA confirmed suppression of IFN-γ response, TNF-NF-κB, IL-6 signaling, and EMT; while metabolic and epithelial repair programs were enhanced following treatment (Figure 6*C*-*D*). Immunofluorescence confirmed significant reductions in p-STAT1, CXCL9, MPO, S100A8/9, and Vimentin in fistula tracts following anti-IFN-γ treatment (Figure 6*E*). The data demonstrate that IFN-γ pathway blockade reverses the transcriptomic and molecular features of PFCD, leading to fistula healing.

**Figure 6.**
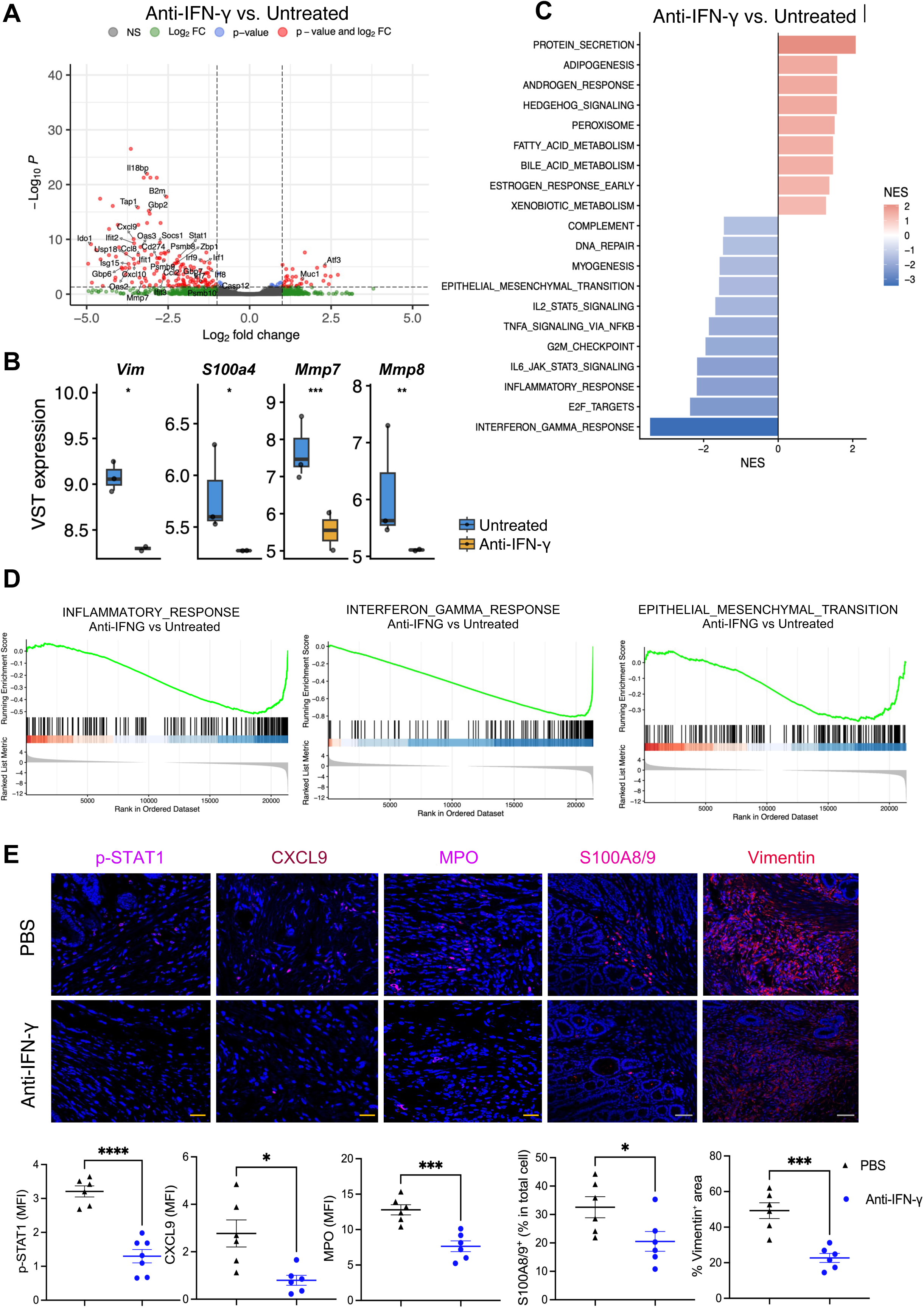
Transcriptomic and protein-level characterization of fistula tissues following anti-IFN-γ treatment in PFCD mouse models. (*A*) Volcano plot of RNA-seq data of fistula tissues from anti-IFN-γ treated vs. untreated WT-TNBS model. (*B*) Normalized expression of *Vim*, *S100a4*, *Mmp7*, and *Mmp8* in fistula tissues of anti-IFN-γ treated vs. untreated WT-TNBS model (n=3). (*C*) Bar chart of NES scores from GSEA comparing anti-IFN-γ treated vs. untreated fistula tissues. (*D*) GSEA plots of inflammatory response, IFN-γ response, and EMT gene sets following anti-IFN-γ treatment. (*E*) Representative immunofluorescence images and quantification of p-STAT1, CXCL9, MPO, S100A8/9, and Vimentin in fistula tracts of anti-IFN-γ treated vs. untreated mice (n=6). Scale bars, 20 µm (p-STAT1, CXCL9, MPO) and 50 µm (S100A8/9, Vimentin). MFI, mean fluorescence intensity. Data are presented as mean ± SEM. Statistical comparisons were performed using unpaired t-test. Experiments were repeated at least 3 times. \**p*<0.05, \*\**p*<0.01, \*\*\**p*<0.001, \*\*\*\**p*<0.0001.

### Co-targeting IFN-γ and TNF-α augments therapeutic efficacy in PFCD models

Although anti-TNF remains a mainstay of PFCD treatment, most patients fail to achieve adequate and long-term response^1^, highlighting the need for more effective therapeutic strategies. Given the vital impact of IFN-γ pathway in driving PFCD as shown above, we next investigated whether combining anti-IFN-γ with anti-TNF-α treatment could achieve superior therapeutic outcomes. The combined treatment achieved significantly more reductions in MAGNIFI-CD scores compared to anti-TNF-α monotherapy at Week 1 and 2 in both WT-TNBS and *Il10^−/−^* models (Figure 7*A*). Mechanistically, combination therapy led to greater suppression of inflammatory markers MPO and S100A8/9, and tissue remodeling marker Vimentin versus anti-TNF-α alone (Figure 7*B*). Thus, co-targeting IFN-γ and TNF-α may offer a better therapeutic strategy for patients with refractory PFCD.

**Figure 7.**
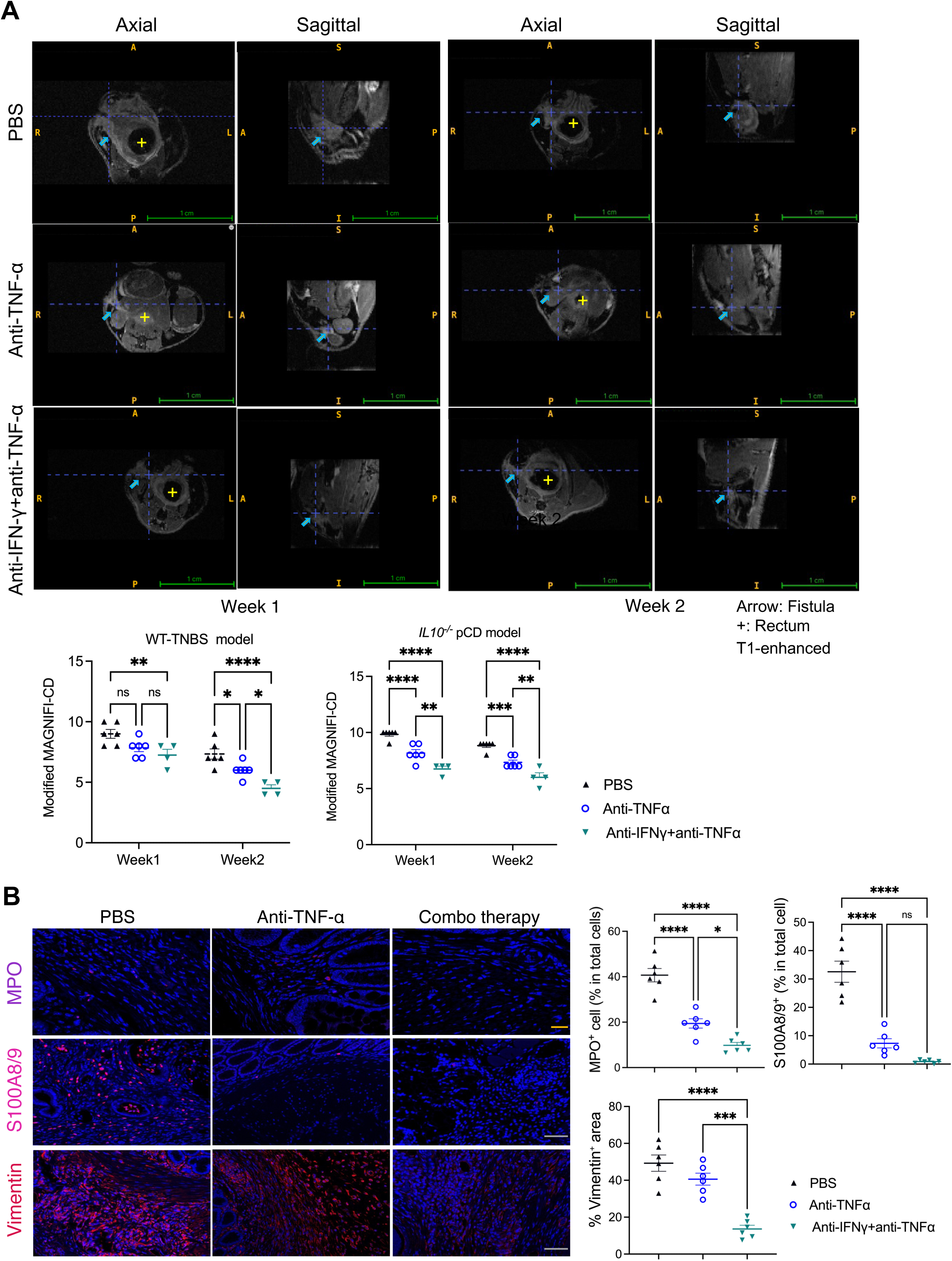
Combined IFN-γ and TNF-α blockade demonstrates superior benefits for fistula healing compared to anti-TNF monotherapy in PFCD mouse models. (A) Representative T1-weighted with contrast axial and sagittal MRI images at Week 1 and 2 in WT-TNBS and *Il10^−/−^* models treated with anti-TNF-α or anti-IFN-γ combined with anti-TNF-α. Modified MAGNIFI-CD scores at Week 1 and 2 across treatment groups (n=4-6). Arrow: fistula tract; +: rectum. (*B*) Representative immunofluorescence images and quantification of MPO, S100A8/9, and Vimentin in fistula tracts from the above treatment groups (n=6). Scale bars, 20 µm (MPO) and 50 µm (S100A8/9, Vimentin). Data are presented as mean ± SEM. Statistical comparisons were performed using one-way ANOVA with Tukey’s post-hoc test. Experiments were repeated at least 3 times. \**p*<0.05, \*\**p*<0.01, \*\*\**p*<0.001, \*\*\*\**p*<0.0001.

## Discussion

PFCD is a highly complex disease likely arising from an intricate interplay of genetic susceptibility, environmental triggers, immune dysregulation, mucosal barrier defects, and architectural tissue stress^4, 24^. Mechanistic studies and therapeutic development in PFCD have been limited by the lack of reliable and physiologically relevant preclinical models. Larger animals are constrained by a lack of relevance to human disease, high costs, low throughput, and difficulty for genetic manipulation. To address this critical gap and unmet need, we established three mouse models of PFCD in which chronic, inflammatory perianal fistulas were created in the setting of CD-like intestinal inflammation. A major strength of our study lies in the deliberate selection of the *Il10*^−/−^ and *TNF^Δ69AU/+^* genetic lines alongside the WT-TNBS chemical-induced model of CD. The *Il10*^−/−^ model recapitulates key features of CD, including transmural inflammation and Th1/Th17-mediated immune responses^25^. The *TNF^Δ69AU/+^*heterozygous model, which lacks macroscopic intestinal inflammation before 6 months of age^16^, does develop proctitis on histology following perianal fistula creation.

We identified shared features of PFCD in all our models that closely resemble findings in PFCD patients by comparing histopathology, fistula tissue microenvironment, and transcriptomic characteristics with non-IBD fistulas. Gene expression analysis identified consistent upregulation of inflammatory and tissue remodeling pathways that have been reported in human PFCD, including IL-17^26^, TNF-α^27^, TL1A-DR3^28^ and IFN-γ^5^ signaling, alongside EMT^7^, fibrosis^29^, and type 2 immune activation^30^. Microbiome profiling further indicated a consistent pattern of dysbiosis across PFCD models. *Alistipes*, a genus negatively associated with IBD and known for production of anti-inflammatory metabolite, was significantly reduced in the models^19, 31^. In contrast, the PFCD models enriched *Muribaculum*, which has been reported to produce the cardiolipin MiCL-1 that induces pro-inflammatory cytokines including TNF-α and IL-6^32^. The convergence of these signatures across PFCD models with pathways reported in human PFCD strongly supports their translational fidelity. In addition, the PFCD models are amenable to non-invasive and quantitative assessment by MRI and therapeutically responsive to anti-TNF and upadacitinib, two current treatments for PFCD patients. This versatile platform provides a powerful and scalable framework for genetic dissection of disease mechanisms and preclinical drug screening.

The role of IFN-γ in PFCD pathogenesis has not been well understood. Our data demonstrate consistent IFN-γ pathway activation across all three PFCD mouse models and human PFCD, with shared producers of IFN-γ including CD8⁺ T cells, NK cells, and ILC1s. We also uncovered that IFN-γ not only drives inflammation but also plays a crucial role in pathogenic tissue remodeling in PFCD. EMT has emerged as a key mechanism in CD-associated fistula formation^6, 7, 33^. Under chronic inflammatory conditions, intestinal epithelial cells may undergo EMT in response to immunological triggers or bacterial invasion, losing their epithelial polarity and cell-cell adhesion while acquiring mesenchymal traits including increased migratory and invasive capacity^7^. Our transcriptomic analyses identified IFN-γ signaling as an essential upstream activator of mesenchymal reprogramming in PFCD. As IFN-γ expression showed strong positive correlations with a broad set of EMT-related genes in human PFCD fistulas, this association was largely absent in IPF tissues. While IFN-γ has been reported to promote EMT in tumor contexts through multiple mechanisms, including activation of JAK-STAT1 signaling and PD-L1, its role in PFCD has not been previously characterized. Transitional cells, which line the non-epithelialized areas of CD fistulas and may sustain/promote fistula tract through an aberrant EMT program driven by TGF-β, TNF, and IL-13^30^, and FAS fibroblasts^22^, which are implicated in ECM remodeling, invasion, fibrosis, and immune cell recruitment, may represent two key cellular constituents of fistula establishment, persistence, and progression in PFCD. The selective elevation of IFN-γ response scores in both cell populations raises the possibility that IFN-γ serves as a previously unrecognized signal stimulating these fistula-promoting cells. Spatial transcriptomic analysis of two independent cohorts further supports this relationship, by reflecting a spatially organized program maintained by immune-epithelial and immune-stromal cell interactions that are unique to PFCD.

One of the most significant findings in our study is the therapeutic efficacy of IFN-γ pathway antagonists in reversing inflammatory and pathogenic tissue-remodeling programs and accelerating fistula healing. In an early clinical trial for luminal CD, systemic IFN-γ neutralization with fontolizumab was well tolerated but produced only modest and inconsistent clinical benefits^34^. It is worth noting that anti-IFN-γ therapy was never tested in fistulizing disease specifically. Here, our results indicate that IFN-γ may play a unique fistula-genic role in the PFCD context, not only as a culprit of mucosal inflammation but as a central regulator of the structural remodeling programs that sustain/promote fistulizing disease. Consistent with this, blockade of IFN-γ and its receptor IFNGR1 promptly improved PFCD by attenuating inflammatory and EMT markers in our mouse models. These preclinical findings, together with the evidence of a disease-specific IFN-γ-EMT axis in patient samples, may help bring interest in IFN-γ-targeted therapy for PFCD. Furthermore, blocking IFN-γ and TNF-α simultaneously may be a viable option for PFCD patients refractory to current treatment.

Several limitations should be acknowledged. First, although our models reproduce many key clinical, histologic, and molecular features of human PFCD, they are initiated by surgical creation of a fistula tract rather than arising spontaneously. Therefore, these models primarily capture mechanisms of fistula persistence and healing rather than the earliest event in fistula initiation. Second, although our transcriptomic, spatial, and pharmacologic data strongly suggest IFN-γ as a key driver of fistula-associated tissue remodeling, the precise mechanism remains to be defined. We expect that our mouse models will facilitate future studies using cell type-specific knockout and lineage-tracing approaches to dissect the cellular circuitry underlying PFCD pathogenesis and therapeutic response. Finally, while IFN-γ signaling is hyperactivated in human PFCD and anti-IFN-γ therapy showed robust efficacy in our mouse models, direct clinical evidence supporting IFN-γ blockade in patients with PFCD remains limited. Future clinical studies are needed to confirm the translational relevance of these findings.

To summarize, we have established three mouse models with chronic and inflammatory perianal fistulas that closely resemble human PFCD based on clinical examination, MRI, histology/immunostaining, and molecular and cellular analyses. The models sustain patent fistula tracts for at least 5 weeks after wire removal, making them reliable tools for mechanistic studies and preclinical therapeutic testing. To the best of our knowledge, these are the first reliable, consistent, clinically relevant, and therapeutically responsive mouse models for PFCD. Importantly, the prompt improvement of mouse fistulas following anti-IFN-γ/IFNGR1 treatment supports the IFN-γ pathway as a novel therapeutic target for PFCD.

## Conflict-of-interest statement

David H. Ballard, MD reports consulting for Direct Biologics. Parakkal Deepak, MBBS MS received research support under a sponsored research agreement unrelated to the data in the study and/or consulting from AbbVie, Arena Pharmaceuticals, Boehringer Ingelheim, Bristol Myers Squibb, Janssen, Pfizer, Prometheus Biosciences, Takeda Pharmaceuticals, Roche Genentech, Scipher Medicine, Fresenius Kabi, Teva Pharmaceuticals, Landos Pharmaceuticals, Iterative scopes and CorEvitas, LLC.

## Acknowledgments

We thank the Washington University (WU) Small Animal Magnetic Resonance Facility (S10OD026913) for help with MRI and the Genome Access Technology Center for RNA sequencing. This study was supported by an internal grant provided to Siyan Cao and a WU Mallinckrodt Institute of Radiology Pilot Fund (Siyan Cao). Siyan Cao was also supported by an NIH/NIDDK K08 Clinical Investigator Award, Crohn’s & Colitis Foundation (CCF) Career Development Award, American Gastroenterological Association Pilot Research Awards, American College of Gastroenterology (ACG) Clinical Research Award, the Lawrence C. Pakula, MD IBD Education & Innovation Fund, the Doris Duke COVID-19 Fund to Retain Clinical Scientists Program, DDRCC Pilot and Feasibility Award, WU Clinical and Translational Research Funding Program and Precision Health Innovation Award, and research funding from Drury Hotels Company, LLC. David H. Ballard reports research funding from the Radiological Society of North America, the CCF, and the Helmsley Charitable Trust. Parakkal Deepak was supported by an ACG Junior Faculty Development Award, IBD Plexus of the CCF, and Leo & Carean Goss Crohn’s Disease Research Fund. Gwendalyn J. Randolph was supported by NIH R01 AI168044.

## CRediT Authorship Contributions

Xin Yao, MD (Data curation: Lead; Formal analysis: Lead; Investigation: Lead; Methodology: Lead; Writing – original draft: Lead; Writing – review & editing: Equal)

Kaiming Ma, BA (Data curation: Equal; Formal analysis: Equal; Investigation: Equal; Methodology: Equal; Writing – original draft: Lead)

David H. Ballard, MD (Formal analysis: Equal; Methodology: Equal)

Echo Zhu, PhD (Data curation: Equal; Formal analysis: Equal)

Xiuli Liu, MD, PhD (Formal analysis: Supporting; Methodology: Equal)

Lingjie Huang, MD, PhD (Data curation: Supporting)

Chuwen Tian, MD (Data curation: Supporting)

James D. Quirk, PhD (Methodology: Supporting)

Heather S. Ruiz (Methodology: Supporting)

Tingyi Tan, MS (Formal analysis: Supporting)

Matthew A. Ciorba, MD (Investigation: Supporting)

Gwendalyn J. Randolph, PhD (Resource: Supporting)

Parakkal Deepak, MBBS, MS (Investigation: Supporting)

Siyan Cao, MD, PhD (Conceptualization: Lead; Funding acquisition: Lead; Data curation: Lead; Formal analysis: Lead; Investigation: Lead; Methodology: Lead; Resource: Lead; Project administration: Lead; Writing –original draft: Lead; Writing – review & editing: Lead)

## Data Transparency Statement

Original data and materials are available from the authors upon reasonable request.

## Notes

### Competing Interest Statement

pending patent for the mouse models described in this study

