## Supplemental for "Novel mouse models for perianal fistulizing Crohn’s disease reveal therapeutic value of interferon-γ antagonists"

Xin Yao, et al.

### Supplemental Methods

#### MRI for PFCD mouse models

The mice were anesthetized using 3% isoflurane and maintained at 1%-1.5% for the duration of the scanning. Mice were injected subcutaneously with 300 µl of a 1:10 dilution of Dotarem (gadoterate meglumine; Guerbet, Princeton, NJ) and positioned supine in a Bruker Biospec 9.4-Tesla MRI (Ettlingen, Germany) using a 4-channel 'rat brain' array receiver coil paired with an 86 mm ID volume transmitter. Contrast-enhanced T1-weighted 3D FLASH images were acquired axially at 125 µm isotropic resolution, with the following parameters: repetition time = 15 ms, echo time = 2.5 ms, FOV = 12x24x12 mm, flip angle = 10 degrees, 3 averages.

#### Drug administration

All treatments were administered via intraperitoneal (IP) injection over a 2-week treatment period. For anti-TNF-α treatment, InVivoMAb anti-mouse TNF antibody (Bio X Cell, Catalog #BE0058) was administered at 0.5 mg/mouse twice weekly. For anti-IFN-γ treatment, InVivoMAb anti-mouse IFN-γ antibody (Bio X Cell, Catalog #BE0055, Clone XMG1.2, Rat IgG1, κ) was administered at 0.5-1 mg/mouse twice weekly. Anti-IFNGR1 (Bio X Cell, Catalog #BE0029, Clone GR-20, Rat IgG2a) antibody was similarly administered at 250 µg/mouse twice weekly, with PBS as the vehicle control for these treatments. For combination therapy, mice received concurrent IP injections of anti-IFN-γ (1 mg/mouse) and anti-TNF-α (0.5 mg/mouse) twice weekly. Upadacitinib (JAK1-selective inhibitor, Bio-Techne, Catalog #7783) was dissolved in DMSO and administered at 20 mg/kg body weight every other day, with DMSO-treated animals serving as vehicle controls.

**Generation and analysis of bulk RNA-seq data**

The entire perianal fistula tracts were dissected and harvested from the models and snap-frozen for batched RNA isolation. Tissues were minced into small pieces in RLT lysis buffer supplemented with 10% β-mercaptoethanol and vortexed for 3 minutes with stainless steel beads. Total RNA was then extracted using the QIAGEN RNeasy Plus Micro Kit (Qiagen, 74034) according to the manufacturer's instructions. RNA integrity was assessed by determining the RNA integrity number (RIN) using the Agilent Bioanalyzer (Agilent Technologies). cDNA was prepared using the SMARTer Ultra Low RNA kit for Illumina Sequencing (Takara-Clontech) at the Genome Technology Access Center (GTAC) at Washington University as recently described by Cao et al[1]. Aligned gene counts were processed using the DESeq2 package (v1.44.0) with R (version 4.3.2). Genes with fewer than 10 counts among all samples were excluded. For analysis of differentially expressed genes (DEGs), only protein-coding genes with an average expression of more than 100 counts per sample were used. The threshold of DEGs was set at an adjusted P value of less than 0.05 and a |log2 fold change| >1. Gene set enrichment analysis (GSEA) was performed using the clusterProfiler package (v4.12.6) with R. Genes for GSEA were filtered as protein-coding genes with an average expression of more than 100 counts and an adjusted P value of less than 0.1.

**Reanalysis of published datasets from Cao et al. and McGregor et al.**

Single-cell RNA sequencing and spatial transcriptomic data from human PFCD fistula tissue were obtained from Cao et al. (2025). Cell type annotations were applied as originally described. IFN-γ signaling and epithelial-mesenchymal transition (EMT) module scores were calculated using the GOBP_RESPONSE_TO_TYPE_II_INTERFERON and GOBP_EPITHELIAL_TO_MESENCHYMAL_TRANSITION gene sets from MSigDB (msigdbr v26.1.0) with AddModuleScore in Seurat (v5.4.0). Pseudo-bulk correlation between IFN-γ signaling and EMT scores was assessed across cell types in both idiopathic fistula (IPF) and PFCD cohorts.

Published spatial transcriptomic data from McGregor et al. (2026) were obtained and loaded into Seurat. Module scores for IFN-γ signaling and EMT were calculated for each Visium spot using the AddModuleScore function in Seurat (v5.4.0), based on the SCT-normalized assay. For spatial visualization of IFN-$\gamma^+$ dominant immune cells, additional module scores were calculated for $CD8^+$ T cells (CD8A, CD8B, GZMB, GZMK, PRF1, IFNG, CXCR3) and ILC1/NK cells (NKG7, KLRD1, FCGR3A, NCAM1, FGFBP2, IL7R, TBX21, CXCR3, KIT, EOMES). A combined IFN-γ/EMT score was computed by summing the z-scaled IFN-γ and EMT module scores for each spot. For each spot, the dominant immune cell type was assigned based on the higher of the two module scores, provided the score exceeded a threshold of 0.1; spots below this threshold were excluded from visualization. Spatial plots were generated using ggplot2 (v3.5.0) in R, with the combined IFN-γ/EMT score. Co-expression analysis was performed on spatial transcriptomics data from Visium slides (4 diverticular fistula and 8 perianal fistula). Spots were classified as co-expressing if both IFN-γ score and EMT score exceeded 0. The percentage of co-expressing spots was calculated for each slide individually. To account for the non-independence of spots within the same slide, a generalized linear mixed model (GLMM) was fitted using the glmer function from the lme4 package (v1.1.37) in R (v4.4.0), with co-expression status (binary) as the outcome, disease group as a fixed effect, and slide ID as a random effect (binomial family, logit link). Statistical significance was determined by the Wald z-test as implemented in the lmerTest package (v3.1.3). Bar plots display the mean percentage of co-expressing spots per group, with error bars representing the standard error of the mean across slides, and individual slide-level values shown as overlaid points.

**Flow cytometry**

Perianal fistulas were dissected from the models and washed with ice-cold HBSS/HEPES. The tissues were cut into small pieces and digested in complete RPMI containing collagenase IV (Sigma, C5318) at 37°C for 50 min with shaking at 250 RPM. The resulting single-cell suspension was filtered through a 100-μm cell strainer and washed with ice-cold HBSS/HEPES. The following antibodies were used for flow cytometric analysis: CD45 (30-F11, BioLegend), CD3 (17A2, BioLegend), CD19 (6D5, BioLegend), Gr-1 (RB6-8C5, BD Biosciences), NK1.1 (PK136, BioLegend), CD11b (M1/70, BioLegend), CD11c (N418, Thermo Fisher Scientific), F4/80 (BM8, BioLegend), Ly6C (HK1.4.rMAb, BioLegend), Ly6G (1A8, BD Biosciences), CD4 (RM4-5, BioLegend), CD8a (53-6.7, BioLegend), CD90.2 (30-H12, BioLegend), RORγt (AFKJS-9, Thermo Fisher Scientific), FoxP3 (MF-14, BioLegend), T-bet (eBio4B10, Thermo Fisher Scientific), TNF-α (MP6-XT22, BioLegend), IFN-γ (XMG1.2, BioLegend). Following surface marker staining, cells were fixed and permeabilized using the Foxp3/Transcription Factor Staining Buffer Set (Invitrogen, cat. 00-5523-00), followed by staining for intracellular targets. Stained cells were analyzed on a FACSymphony A3 cytometer (BD Biosciences). Analysis was performed using FlowJo, version 10.8.1 (FlowJo LLC).

**Immunofluorescence**

Unstained sections were cut from formalin-fixed, paraffin-embedded tissue blocks. Slides were deparaffinized by three changes of xylene, followed by rehydration through three changes of isopropanol, each for 5 minutes. Sections were then washed by distilled water for 15 minutes. Heat-induced antigen retrieval was performed by boiling the slides for 30 minutes in the citrate buffer (pH6.0; Sigma, C9999). After washing three times for 5 minutes in PBS containing 0.075% Tween-20 (PBST), sections were blocked with 2% bovine serum albumin (BSA) and 5% goat serum in PBS for 2 hours at room temperature. Slides were then incubated overnight at 4 °C with primary antibodies diluted in the same blocking buffer. On the following day, sections were washed three times with PBST and incubated with appropriate fluorescently labeled secondary antibodies diluted in 2% BSA for 1 hour at room temperature in the dark. Nuclei were counterstained with DAPI (Thermo Fisher Scientific, 33258) for 20 minutes followed by 3 times of washes. Finally, slides were mounted using ProLong Gold Antifade Mountant (Invitrogen, P36934). Images were acquired using a Zeiss LSM 880 confocal microscope (Carl Zeiss, Germany). Each dot represented one biological sample. The mean fluorescence intensities (MFIs), quantified by ImageJ, within a sample were used for statistical analysis. The same area of regions of interest (ROIs) was randomly chosen from three to four fields per slide. For p-STAT1, nuclear translocation was quantified as the nuclear-to-cytoplasmic (N/C) ratio of mean p-STAT1 fluorescence intensity, with nuclear regions defined by the DAPI channel and cytoplasmic regions defined as the surrounding perinuclear area. IFN-$\gamma^{+}$CD45$^{+}$ double-positive cells and MPO$^{+}$ cells were counted and normalized to the total number of DAPI$^{+}$ nuclei. %CD45$^{+}$, α-SMA, and vimentin area were calculated as positive staining area divided by total tissue area. Measurements were performed blindly to group identity. Data were obtained from at least three mice per group. The following primary antibodies were used for immunofluorescence staining with dilutions: pan-cytokeratin (Thermo Fisher Scientific, MA5-13203, 1:150), CD45 (Thermo Fisher Scientific, 14-0451-82, 1:100), α-SMA (Thermo Fisher Scientific, MA5-44355, 1:150), CD4 (Thermo Fisher Scientific, MA1-146, 1:75), F4/80 (Thermo Fisher Scientific, MA5-16363, 1:50), TNF-α (Thermo Fisher Scientific, 14-7321-81, 1:50), myeloperoxidase (MPO; Bioss, BS-4943R, 1:75), IFN-γ (Bioss, BS-0480R, 1:100), CXCL9 (Invitrogen, PI701117, 1:100), phospho-JAK2 (Invitrogen, PIMA542424, 1:100), vimentin (Abcam, ab92547, 1:150), S100A8/S100A9 (Abcam, ab288715, 1:150), and phospho-STAT1 (Cell Signaling Technology, 44-376G, 1:100). Secondary antibodies included Alexa Fluor–conjugated goat anti-rat IgG (Invitrogen, A21247, 1:100), goat anti-mouse IgG (Invitrogen, A11001, 1:100), and goat anti-rabbit IgG (Invitrogen, A11011, 1:100).

**Histology score**

The histology of mouse perianal fistulas (H&E staining) was scored by a certified GI pathologist using the following criteria adopted from human PFCD pathology: mild acute inflammation: 1; mild acute and chronic inflammation: 2; moderate chronic and acute inflammation: 3; dense chronic and acute inflammation: 4.

### Statistics

Statistical significance was determined using a 2-tailed Student's t-test, 1-way ANOVA, or 2-way ANOVA, as appropriate, followed by Tukey's multiple-comparisons test for post-hoc analysis. GraphPad Prism 10.0 (GraphPad Software) was used for statistical analysis. A P value of less than 0.05 was considered statistically significant. Data represent the mean ± SEM.

### Study approval

All animal studies were conducted under the approval of the IACUC of Washington University.

### Data availability

We will make data generated in this study, analytic methods, and study materials available to other researchers upon reasonable request.

### Supplemental Table1

Modified MAGNIFI-CD scoring index for murine perianal fistula models.

| MAGNIFI-CD index for PFCD patients | Modified MAGNIFI-CD for mice (0-14) | Rationale |
|---|---|---|
| Number of fistula tracts (0 = none; 1 = single; 2 = complex) | Not included | Not applicable in the murine model. |
| Fistula length (0 = ≤2.5 cm; 1 = >2.5 cm; 2=> 5cm) | Not included | Not included due to limited length variability in the mechanically induced murine model. |
| Fistula tract visibility (Additional item) | Fistula tract visibility (0 = not visible; 1 = <50% visible; 2 = >50% visible) | Included to provide the anatomic confirmation of tract conspicuity. |
| Internal opening (IO) visibility (Additional item) | IO visibility (0 = not visible; 1 = visible) | Included as an anatomic confirmation of the rectal communication. |
| External opening (EO) visibility (Additional item) | EO visibility (0 = not visible; 1 = visible) | Included as an anatomic confirmation of the perianal skin communication. |
| Hyperintensity of the primary tract on post-contrast T1-weighted images (0=absent; 1=pronounced) | Hyperintensity of the primary tract on post-contrast T1-weighted images (0 = none; 1 = focal/mild; 2 = partial continuous; 3 = extensive/pronounced) | Retained as the primary MRI feature reflecting fistula inflammatory activity; scale adapted for murine imaging. |
| Dominant feature of fistula tract (0 = fibrous; 1 = granulation tissue; 2 = fluid/pus) | Dominant feature of fistula tract (0 = fibrous; 1 = granulation tissue; 2 = fluid/pus) | Retained to categorize dominant tract composition relevant to inflammatory status. |
| Presence of proctitis | Presence of proctitis (0 = absent; 1 = mild; 2 = pronounced) | Retained as the primary MRI feature reflecting rectal inflammatory activity; scale adapted for murine imaging. |
| Extension of fistula (0 = absent; 1 = horseshoe ; supralevatoric) | Not included | Not applicable in the murine model. |
| Inflammatory mass/collections (0 = absent; 1 = focal; 2 = diffuse; 3 = collections small; 4 = collections medium; 5 = collections large) | Inflammatory mass/collections (0 = none; 1 = small <2 mm; 2 = medium 2–3 mm; 3 = large >3 mm) | Modified based on murine size constraints. |

Abbreviations: MAGNIFI-CD, Magnetic Resonance Index for Fistula Imaging in Crohn's Disease; IO, internal opening; EO, external opening; PFCD, perianal fistulizing Crohn's disease; MRI, magnetic resonance imaging.

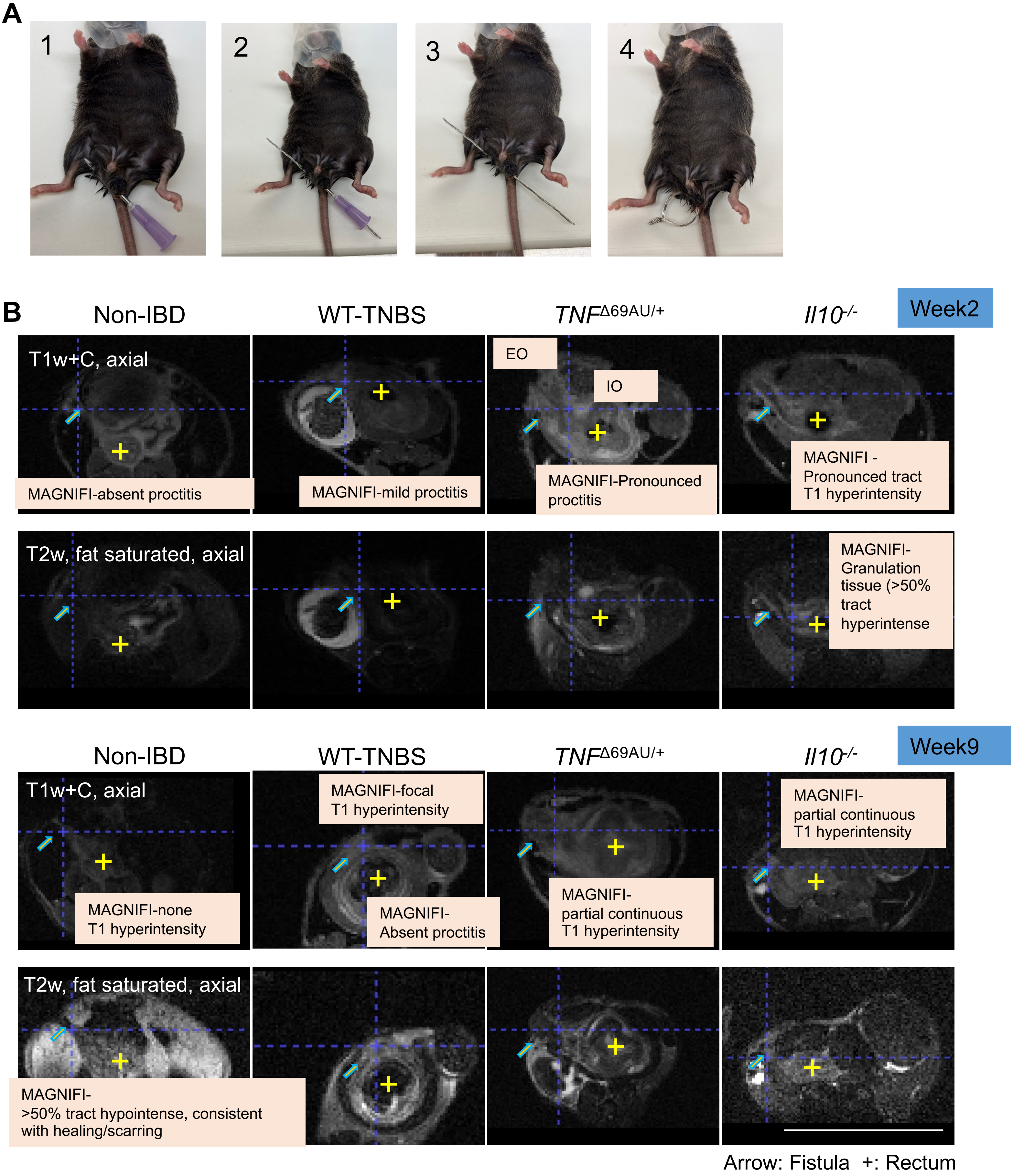


**Supplemental Figure 1. Standardized surgical procedure for perianal fistula induction and MAGNIFI-CD MRI features.** (*A*) The stepwise perianal wire implantation procedure in mice: (1) initial positioning, (2) needle insertion, (3) wire placement, and (4) completion of fistula induction. (*B*) Representative MAGNIFI-CD MRI features at Week 2 and Week 9 across mouse models. Representative T1w+C and T2-weighted fat-saturated axial MRI images at Week 2 and Week 9. Annotated MAGNIFI-CD features are indicated. PFCD mice models demonstrated persistent fistula activity and inflammation compared to Non-IBD mice. Arrow: fistula tract; +: rectum.

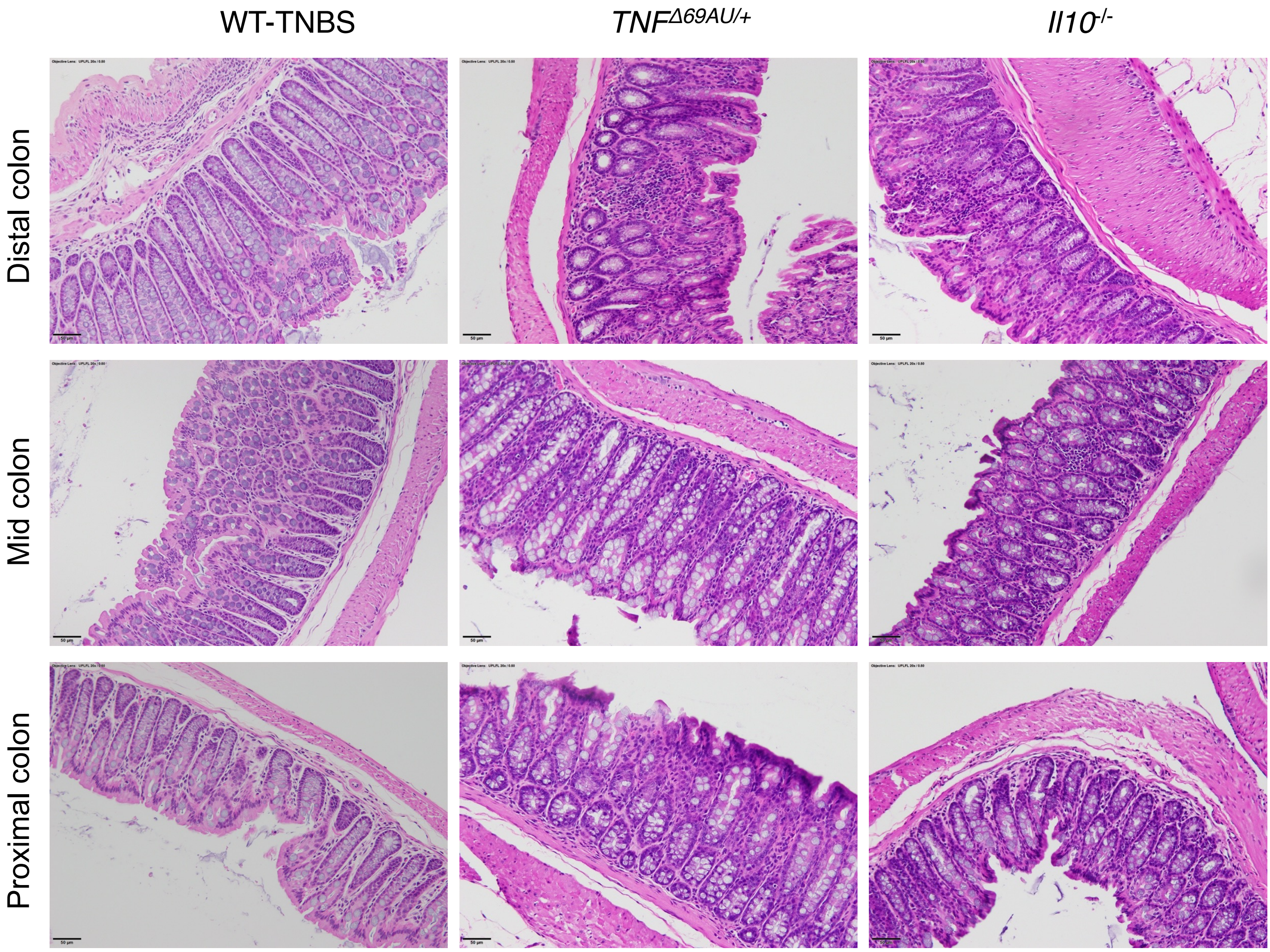


**Supplemental Figure 2. Colon histology of PFCD models.** Representative H&E staining of distal, mid, and proximal colon sections from WT-TNBS, the *TNF*$^{\Delta 69AU/+}$ and *Il10*$^{-/-}$ models.

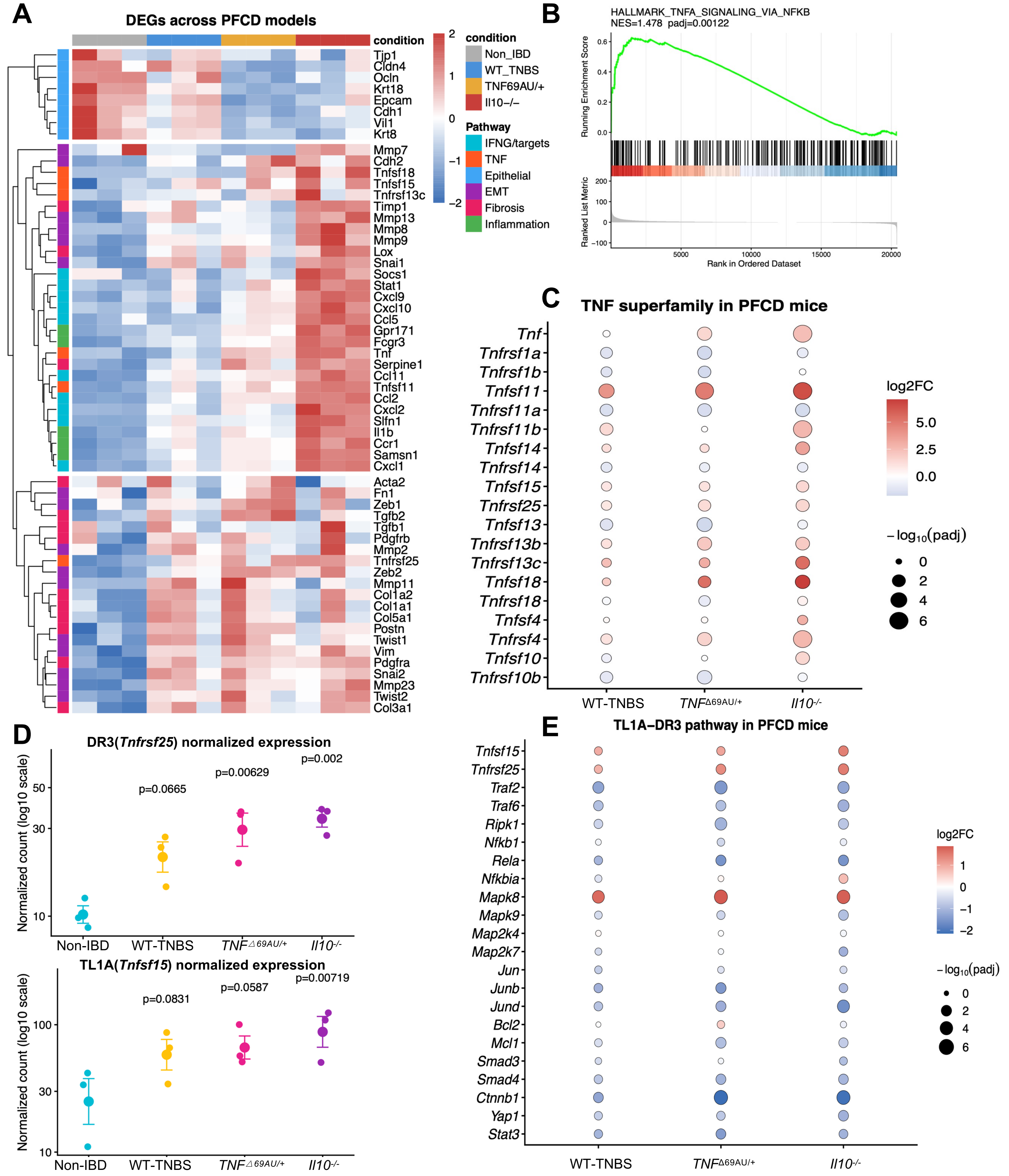


**Supplemental Figure 3. Transcriptomic characterization of shared differentially expressed genes and TL1A-DR3 pathway activation across PFCD models.** (*A*) Heatmap of differentially expressed genes (DEGs) across PFCD models compared to non-IBD controls. Genes are annotated by pathway category. (*B*) Gene Set Enrichment Analysis (GSEA) plot showing enrichment of the HALLMARK_TNFA_SIGNALING_VIA_NFKB gene set in PFCD models. NES = 1.478, *p*= 0.00122. (*C*) Bubble plot showing log2 fold change and statistical significance of TNF superfamily ligand and receptor genes across PFCD models compared to Non-IBD controls. Dot size represents $-\log_{10}$(*padj*) and color indicates log2FC. (*D*) Normalized expression (log10 scale) of DR3 (*Tnfrsf25*, top) and TL1A (*Tnfsf15*, bottom) across non-IBD and PFCD models. P-values from pairwise comparisons are indicated. (*E*) Bubble plot showing TL1A-DR3 pathway core genes across the PFCD models.

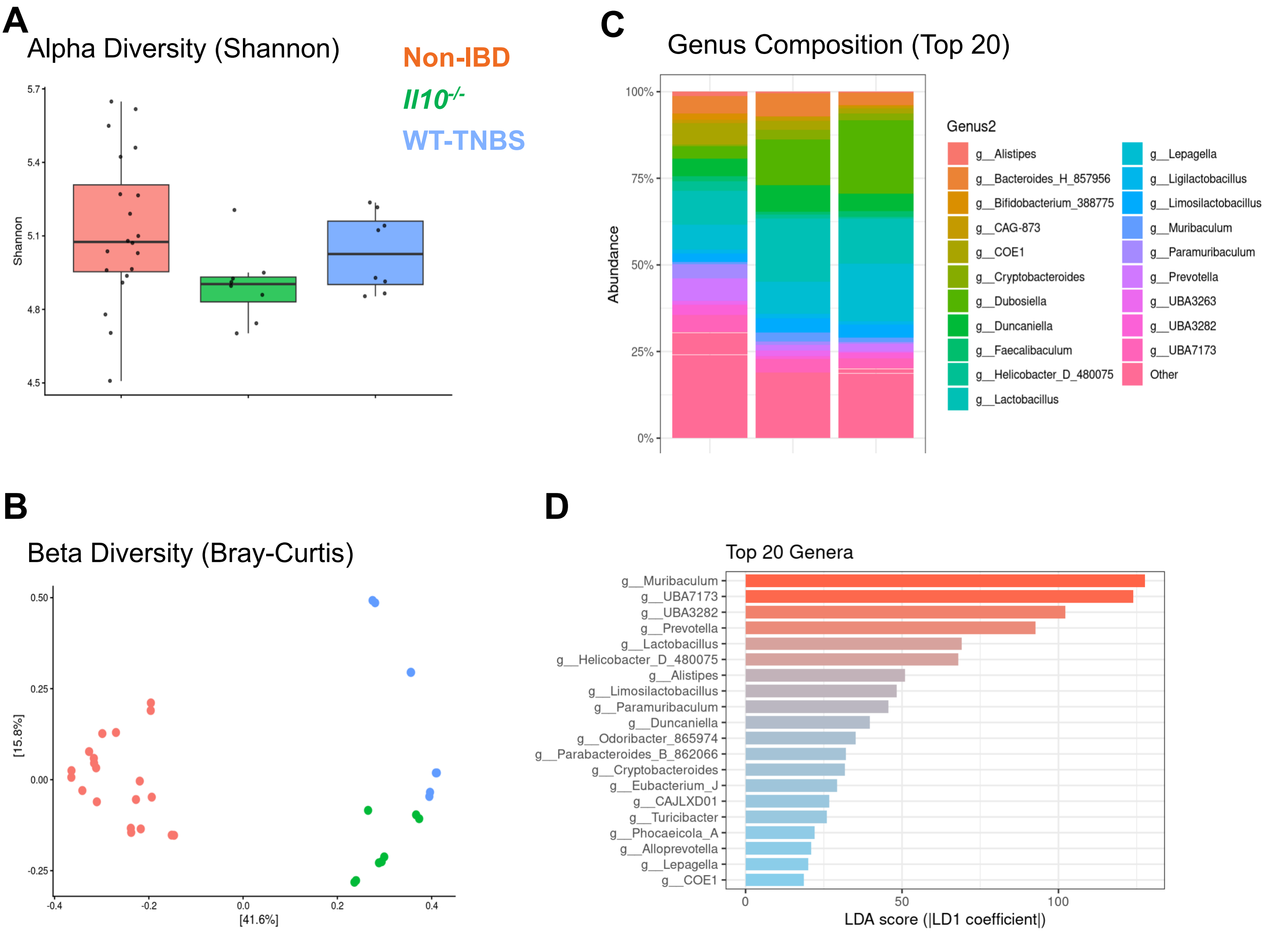


**Supplementary Figure 4. Microbiome profiling and diversity analysis in PFCD models.** (*A*) Alpha diversity measured by the Shannon index across non-IBD fistula, *Il10$^{-/-}$*, and WT-TNBS groups, *p*= 0.06. (*B*) Principal coordinate analysis (PCoA) of beta diversity based on Bray-Curtis dissimilarity among the three groups, *p*= 0.001. (*C*) Stacked bar plots showing the relative abundance of the top 20 bacterial genera. (*D*) Linear discriminant analysis (LDA) effect size (LEfSe) scores (based on LD1 coefficients) identifying the top 20 most differentially abundant genera across the groups.

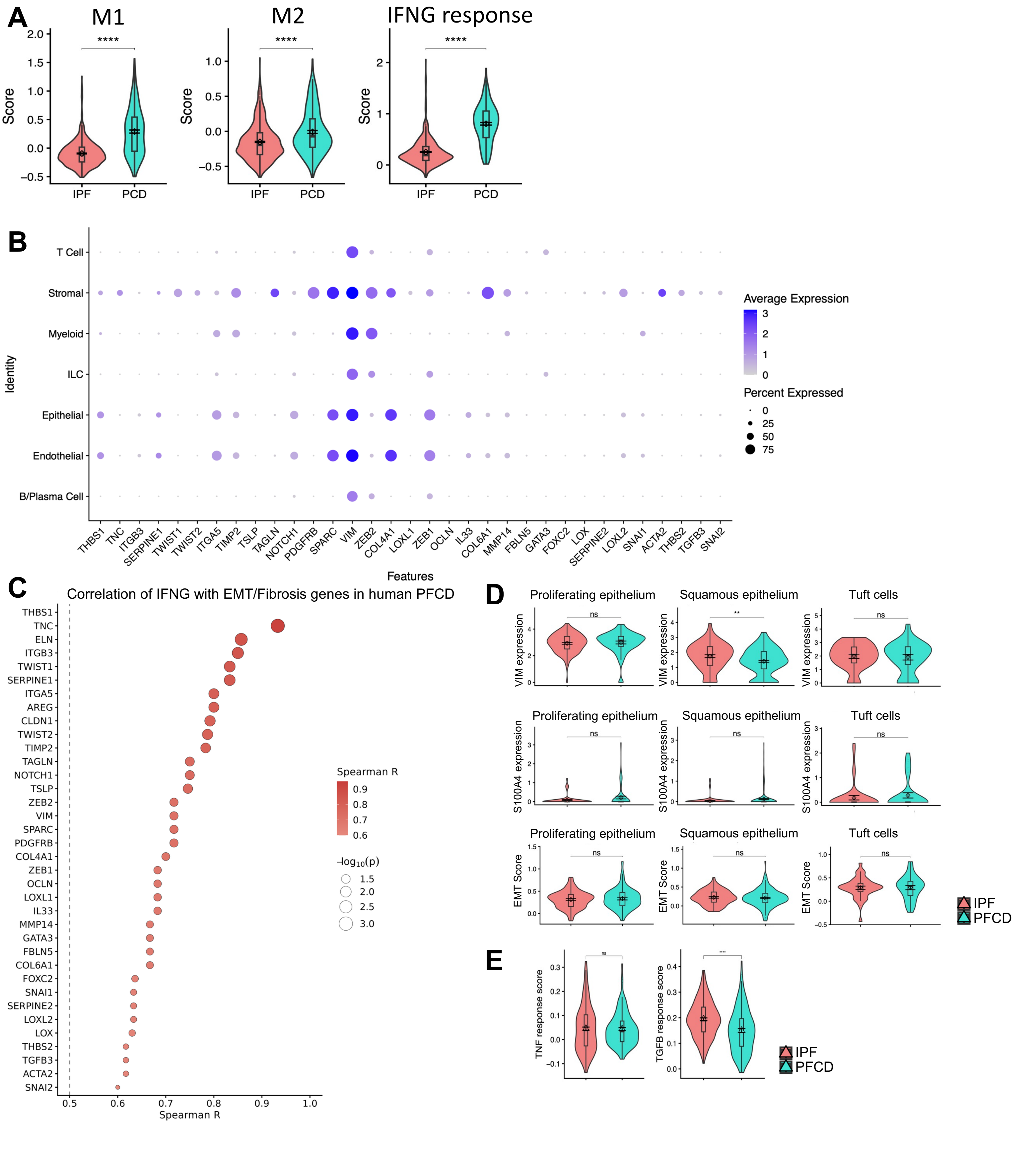


**Supplementary Figure 5. Transcriptomic and pathway analysis of immune activation, EMT, and fibrosis in IPF and PFCD patients.** (*A*) Violin plots comparing macrophage polarization scores and functional activity between IPF and PFCD patients. M1 score (left), M2 score (middle), and IFNG response score (right) are shown for IPF and PFCD groups. (*B*) Dot plot showing average expression and percent expressed of EMT, fibrosis, and tissue remodeling-related genes across major cell type clusters (T Cell, Stromal, Myeloid, ILC, Epithelial, Endothelial, and B/Plasma Cell) in PFCD patient samples. Dot size represents percent of cells expressing the gene and color indicates average expression level. (*C*) Bubble plot showing Spearman correlations between IFNG expression and EMT/fibrosis-related genes in human PFCD. Dot color indicates Spearman R value and dot size represents $-\log_{10}(p)$. **** $p< 0.0001$. (*D*) Violin plots comparing VIM expression (top), S100A4 expression (middle), and EMT score (bottom) in proliferating epithelium, squamous epithelium, and tuft cells between IPF and PFCD patients. ** $p< 0.01$, ns = not significant. (*E*) Violin plots comparing TGF-β response score and TNF response score between IPF and PFCD patients. **** $p< 0.0001$, ns = not significant.

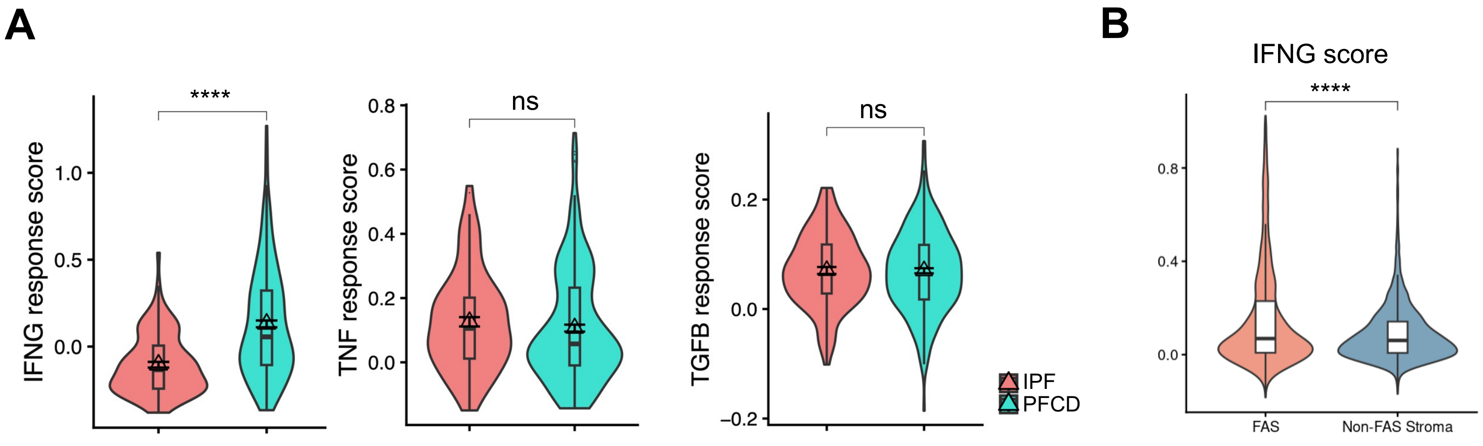


**Supplemental Figure 6. IFN-γ-driven EMT is enhanced in PFCD compared to IPF.** (*A*) Violin plots depicting IFNG, TNF, and TGFB pathway response scores in fistula-associated stromal (FAS) fibroblasts stratified by disease group (IPF vs. PFCD). (*B*) Violin plots comparing IFN-γ scores between FAS fibroblasts and Non-FAS stromal cells. **** $p<0.0001$.

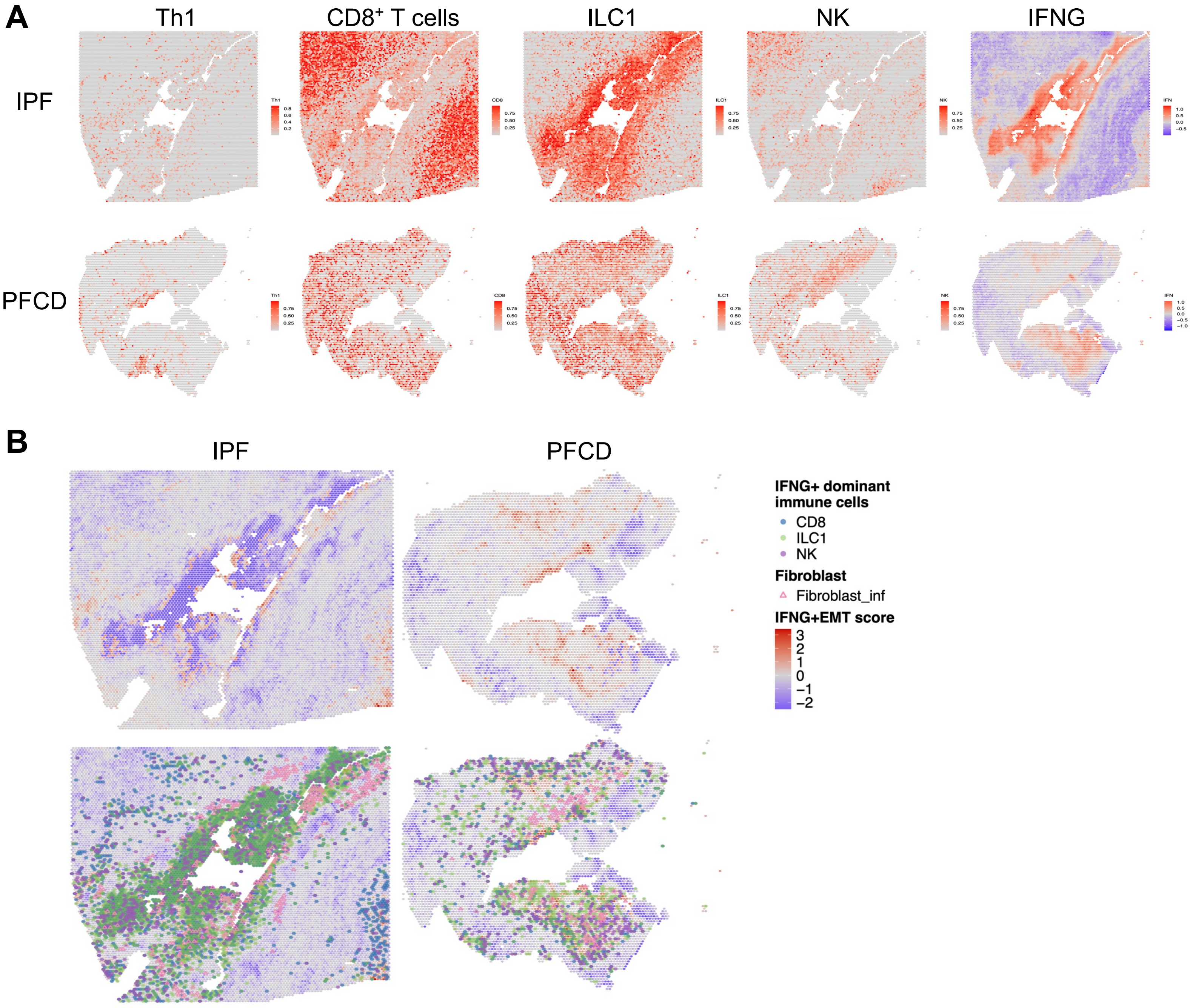


**Supplemental Figure 7. Spatial distribution of IFN-γ$^+$ immune cells, inflammatory fibroblasts, and EMT score in human IPF and PFCD tissues.** (*A*) Spatial transcriptomic maps of Th1 cells, CD8 T cells, ILC1s, NK cells, and IFN-γ score in idiopathic fistula (IPF, upper panels) and PFCD (lower panels). (*B*) Spatial transcriptomic maps showing the combined IFN-γ/EMT score (defined as the sum of z-scaled IFN-γ and EMT module scores) across all tissue spots in IPF and PFCD (upper panels), and the spatial distribution of IFN-γ$^+$ dominant immune cells (CD8$^+$ T cells, ILC1s, and NK cells) and inflammatory fibroblasts (Fibroblast_inf) superimposed on the combined IFN-γ/EMT score (lower panels).

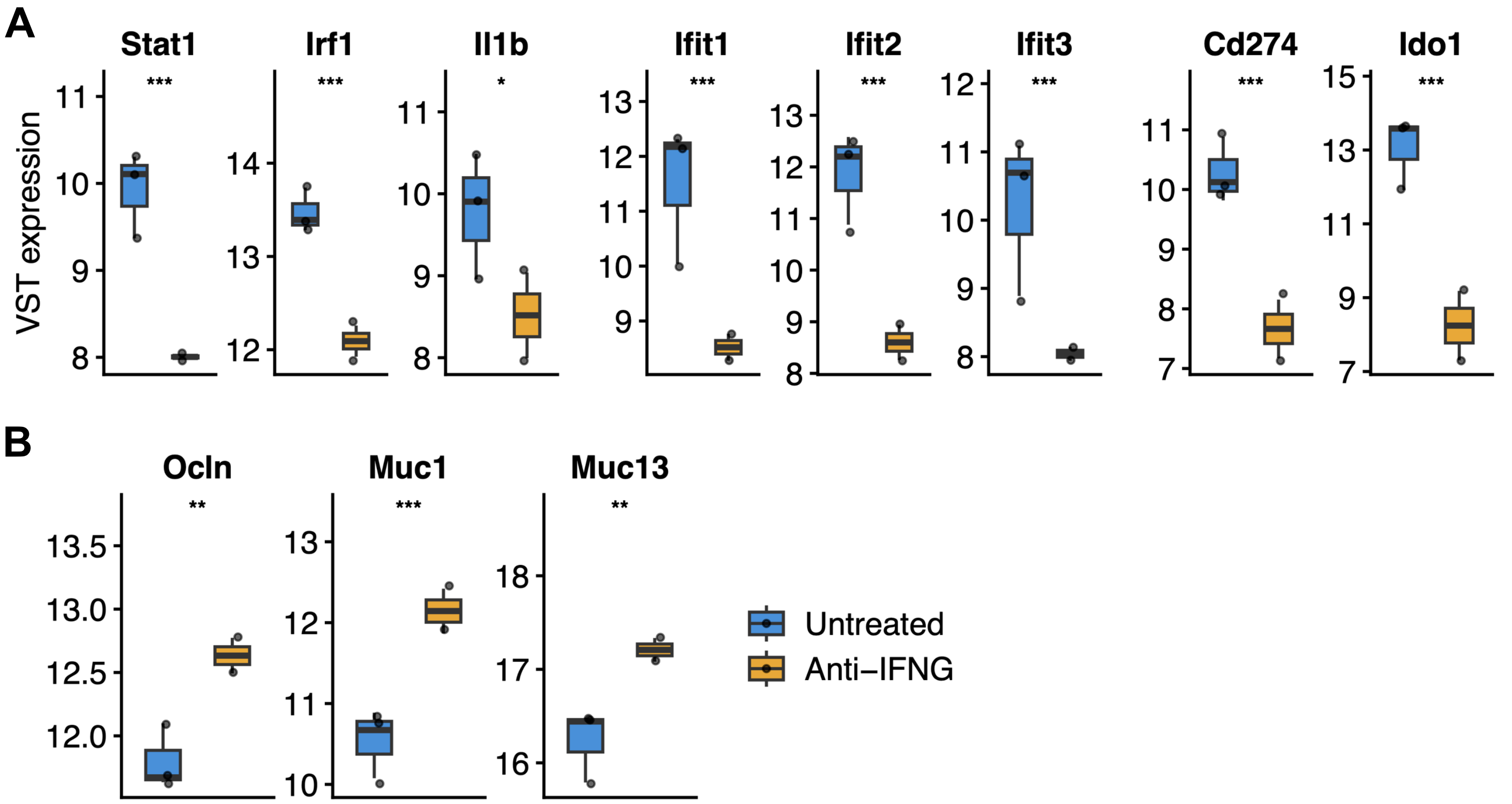


**Supplementary Figure 8. Transcriptomic changes after anti-IFN-γ treatment in PFCD mouse fistula tissue**. (*A*) Box plots showing VST-normalized expression of IFN-γ signaling genes (Stat1, Irf1, Il1b), interferon-stimulated genes (ISG response; Ifit1, Ifit2, Ifit3), and immune suppression markers (Cd274, Ido1) in untreated and anti-IFN-γ-treated fistula tissues. (*B*) Box plots showing expression of epithelial repair genes (Ocln, Muc1, Muc13) in untreated and anti-IFN-γ-treated fistula tissues. Statistical comparisons were performed using Student's t-test. *$p$<0.05, **$p$<0.01, ***$p$<0.001.
